# Prophage induction contributes to alterations in the gut phageome during intestinal inflammation

**DOI:** 10.1101/2024.12.03.626644

**Authors:** Anshul Sinha, Amy Qian, Tommy Boutin, Corinne F. Maurice

## Abstract

Bacteriophages (phages) are abundant members of the gut microbiota and regulators of bacterial communities. During homeostasis, gut phage communities are longitudinally stable and lysogenic replication is dominant. In chronic gut inflammatory disorders, such as inflammatory bowel diseases (IBDs), there are alterations in phage diversity, which may result from changes in phage replication cycle dynamics. Here, we used a combination of *in vitro*, simplified community, and whole-community bioinformatics approaches to investigate whether prophage induction contributes to these alterations. We identified several compounds associated with gut inflammation that induced prophages in commensal gut bacterial isolates. Analysing data from two mouse models of colitis, we observed that shifts in the composition of temperate phages occur over the course of inflammation, supporting a switch from lysogenic to lytic replication. Collectively, our observations support the idea that prophage induction contributes to alterations in the phageome associated with inflammation.

## Introduction

The gut microbiota has a central role in immune homeostasis and is implicated in the development of inflammatory bowel diseases (IBDs), such as Crohn’s Disease (CD) and ulcerative colitis (UC)^1^. IBDs are chronic disorders, characterized by debilitating gastrointestinal symptoms that manifest in periods of flare and remission^2,3^. While these diseases are multifactorial in nature, changes to gut bacterial diversity are hallmarks of disease severity and important to disease progression^4,5^. Alterations to gut bacterial communities in IBDs include reduced diversity of obligate anaerobes belonging to the phyla Bacteroidota and Bacillota^6,7^, which produce metabolites such as short-chain fatty acids that stimulate immunomodulatory responses in the gut^8,9^. In addition, an expansion of invasive facultative anaerobes, mostly Pseudomonadota, contributes to the excess pro-inflammatory responses characteristic of IBDs^6,10^. While these changes to gut bacterial communities are well characterized, the factors responsible for them remain difficult to disentangle. There are drastic changes to the gut microenvironment in CD and UC, including an influx of immune cells to the intestinal epithelium^11^, alterations in host and bacterial-produced metabolites^6,12^, and increased oxygen concentrations in the colon^13^. While these factors may contribute to the observed shifts of bacterial communities during IBDs, the role of other members of the microbiota, such as bacteriophages (phages), in relation to these shifts remains unclear.

Phages are abundant members of the gut microbiota, present at an approximately equal ratio to their bacterial hosts^14,15^. Importantly, phages in the gut shape bacterial diversity^16–18^, alter bacterial fitness^19,20^, and modify the abundances of bacterial-derived metabolites^16^. Through their ability to regulate bacterial communities, phages can also impact mammalian health, as in the case of mouse models of disease (i.e., type 2 diabetes^21^ and IBDs^22,23^) and possibly in the context of human *Clostridioides difficile* infection^24^.

Typically, phages in the gut replicate through the lytic or the lysogenic replication cycle^25,26^. The lytic cycle is characterized by a rapid production of new viral progeny after infection. In the lysogenic cycle, phages integrate into the genome of their bacterial host and passively replicate along with the bacterial chromosome as a prophage^27^. This passive replication continues until a prophage induction event occurs where there is a switch from lysogenic to lytic replication^28,29^. Temperate phages are those capable of undergoing lysogenic and lytic replication, whereas virulent phages only undergo lytic replication. The balance between lytic and lysogenic replication has important consequences on bacterial community dynamics^30^. In healthy adults, lysogenic replication dominates due to high bacterial fitness and density in the gut, which favours phage integration^31,32^. These high rates of lysogenic replication likely allow for phage-bacterial co-existence in the gut and contribute to the longitudinal stability of the microbiota^33^. Still, whether phages and bacteria undergo lytic or lysogenic replication in the gut is highly contextual and dependent on environmental conditions^31,32^, stochastic processes^29^, and on individual phage-bacteria pairs. Upon exposure to certain stressors in the gut, such as medication^34,35^ and certain dietary compounds^36,37^, there is a switch from lysogenic to lytic replication, as prophage induction is closely tied to bacterial stress.

During inflammation, there are shifts to the gut phageome, similar to what is reported for bacterial communities. These changes are characterized by a shift in overall phage diversity^38,39^, an increase in extracellular temperate phages^39^, an enrichment in phages infecting bacteria from the phyla Bacillota^39,40^, high inter-individual variability^41^, and a lack of longitudinal stability^41^. Recent work from our lab further showed that transfer of phages from UC patients to mice with a humanized microbiota exacerbated colitis severity in a mouse model of inflammation^22^. These data imply that changes to phage communities in IBD are not simply a consequence of inflammation but could also contribute to disease progression. Despite these observations of changes to gut phage communities in IBDs, what drives these alterations remains unclear. Clarity on the aetiology of changes to the phageome are critical to understanding gut microbiota dynamics during inflammation.

The observations that extracellular temperate phages are enriched in IBD patients compared to non-IBD controls suggest that prophage induction could be involved in these phage community changes. However, there is a lack of experimental evidence linking the changes in the inflamed gut to prophage induction. Similarly, there is a lack of longitudinal resolution in tracking prophage induction over the course of inflammation. Using a variety of strategies, we assessed whether prophage induction occurs in the context of intestinal inflammation. First, we identified several compounds enriched during inflammation that induce a small collection of gut bacterial prophages *in vitro.* Next, using gnotobiotic mice colonized with the oligo-mouse microbiota 12 (OMM^12^) synthetic bacterial community, we observed evidence of prophage induction in response to experimental colitis. Lastly, we observed shifts in the composition of temperate phages in the context of inflammation in two previously published metagenomic datasets. Taken together, our data add support to the idea that prophage induction contributes to the shifts in phage community composition during intestinal inflammation.

## Results

### Inflammatory compounds inhibit gut bacterial growth and increase VLP production in vitro

During intestinal inflammation, bacterial communities in the gut are exposed to several compounds linked to alterations in the gut microenvironment^6^. To test whether these compounds can induce gut bacterial prophages, we performed an *in vitro* prophage induction assay, screening a set of compounds known to be associated with intestinal inflammation against a small panel of commensal gut bacteria. The bacterial strains *Flavonifractor plautii* YL31, *Bacteroides caecimuris* I48, and *Akkermansia muciniphila* YL44 were selected, as they represent genera common to the mammalian gut microbiota with known inducible prophages^42,43^. A second *A. muciniphila* strain, BAA-835, was selected to determine whether there were strain-specific differences in induction. For each strain, prophages were identified bioinformatically using VIBRANT (Supplemental Table S1). In addition, for *F. plautii* YL31, *B. caecimuris* I48, and *A. muciniphila* YL44, previous studies^42,43^ have identified prophages within these strains that are inducible *in vitro* (Supplemental Table S1). To our knowledge, no studies have tested whether *A. muciniphila* BAA-835 contains inducible prophages; however, using VIBRANT, we bioinformatically predicted 2 prophage regions (Supplemental Table S1). Notably, no prophages were shared between the *A. muciniphila* strains YL44 and BAA*-*835 (Supplemental Table S1).

We screened a panel of 8 compounds on each of these 4 strains, at 2-3 different concentrations, for a total of 68 conditions (Table 1). These compounds broadly belong to 3 classes of compounds known to be enriched in the context of intestinal inflammation: immune cell-derived compounds^44–47^, bile salts^6,12^, and *N*-acyl ethanolamines (*N*AEs)^6,48^. Compounds that had previously been shown to have anti-bacterial activity were prioritized for their inclusion.

**Table 1:** Inflammatory compounds selected for *in vitro* prophage induction assays. Compounds known to be enriched in human IBD were chosen to be screened.

| Compound Class | Compound | Reference | Concentrations Tested |
| --- | --- | --- | --- |
| Immune-derived compounds | Hydrogen peroxide (Thermo-Scientific) | Santhanam (2007)<sup>44</sup>, | 0.25 mM, 1mM |
|  | Peroxynitrite (Millipore-Sigma) | McCafferty (2000)<sup>45</sup>, Kimura (1998)<sup>49</sup> | 5 mM, 10 mM |
|  | Human beta-defensin 3 (hBD-3) (Anaspec) | Fahlgren (2004)<sup>47</sup>, Meisch (2013)<sup>47</sup> | 1 µg/mL, 5 µg/mL |
| | TNF-α (Thermo-Scientific) | Komatsu (2001)<sup>46</sup>, Matsuda (2009)<sup>54</sup> | 10 ng/µL 100 ng/µL |
| Bile salts | Bile salt mix (Oxoid, Thermo-Scientific) | Lloyd-Price (2019)<sup>6</sup>, Franzosa (2019)<sup>12</sup> | 0.125 %, 0.25 % |
|  | Taurocholic acid (Thermo-Scientific) | Lloyd-Price (2019)<sup>6</sup> | 1.25 %, 2.5 % |
| *N*-acyl ethanolamines (NAEs) | Linoleoyl ethanolamide (LEA) (Cayman Chemicals) | Lloyd-Price (2019)<sup>6</sup>, Franzosa (2019)<sup>12</sup> | 0.025 mM, 0.05 mM, 0.1 mM |
|  | Palmitoyl ethanolamide (PEA) (Bio-Techne) | Lloyd-Price (2019)<sup>6</sup>, Franzosa (2019)<sup>12</sup> | 0.05 mM, 0.01 mM |

Similar to Sutcliffe *et al.*,^34^ we used bacterial growth inhibition as an initial screen to identify candidate conditions that could induce prophages *in vitro*. Growth inhibition was defined by a decrease in the area under the bacterial growth curve (AUC) of ≥ 15% in the treatment group compared to the vehicle control^34^. In total, 31/68 (45.58 %) conditions led to a decrease in AUC ≥15 % across the 4 strains (Fig. 1A). In general, bile salts (13/16; 81.25%), reactive oxygen species (ROS) (4/8; 50%) and reactive nitrogen species (RNS) (8/8; 100%) led to particularly high levels of growth inhibition (Fig. 1A). Specifically, peroxynitrite, a RNS found at elevated levels in IBD patients^45,49^, led to growth inhibition across all tested conditions (Fig. 1A). A bile salt mix containing conjugated bile salts, and taurocholic acid, a conjugated bile acid enriched in CD patients^6^, had antibacterial activity in all strains, except *B. caecimuris* (Fig. 1A). *N*AEs, a class of signalling lipid including linoleoyl ethanolamide (LEA) and palmitoylethanolamide (PEA), shown to be elevated in IBD patients^6,12^, only inhibited bacterial growth in 4/20 (20%) conditions tested (Fig. 1A). Interestingly, compared to the other bacterial strains, *B. caecimuris* generally exhibited higher resistance to these inflammatory compounds (5/17; 29.41% growth inhibition) (Fig. 1A). While both *A. muciniphila* strains had similar sensitivity to bile acids and immune-derived compounds, there were strain-specific differences in growth inhibition in response to LEA. These data together suggest that inflammatory compounds can directly inhibit the growth of commensal gut bacteria *in vitro*, with strain and dose-dependent differences in sensitivity.

**Fig. 1.**
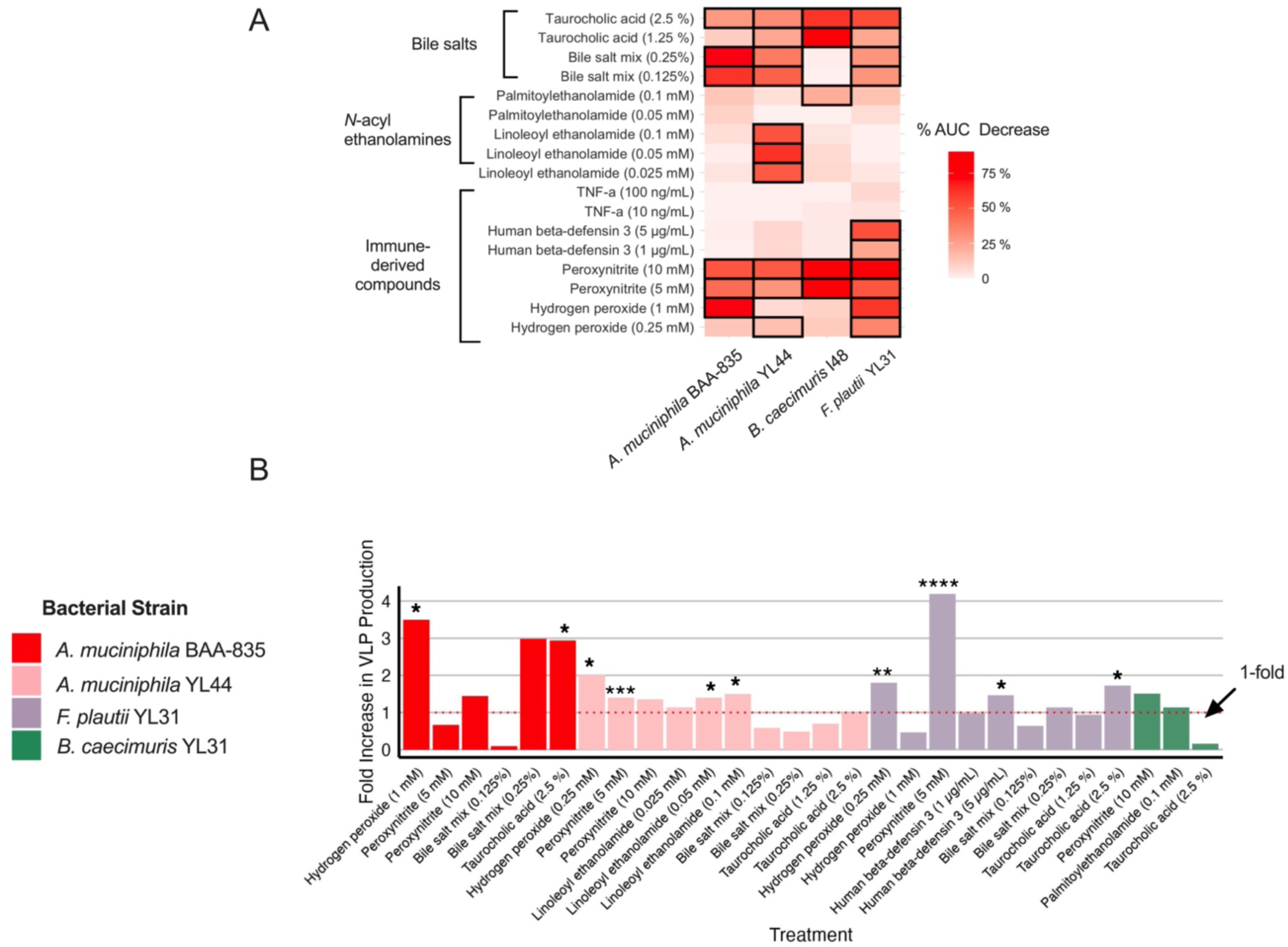
Inflammatory compounds inhibit gut bacterial growth and increase VLP production *in vitro*. A) Heatmap of the percentage decrease in the area under the growth curve (AUC) of bacterial strains exposed to inflammatory compounds compared to vehicle controls. Black boxes represent conditions where the decrease in AUC was ≥ 15%. Bacteria were grown anaerobically in triplicate (n=3) and were treated with inflammatory compounds at the start of growth. Fold-increase in VLP production is relative to the vehicle control. B) VLPs were quantified using epifluorescence microscopy in conditions where the decrease in bacterial growth was 15% ≤ AUC ≤ 75 %. VLPs were quantified from triplicate incubations (n=3), with 2 technical slide replicates. Significant increases in fold-change between treatment and vehicle controls were assessed using an unpaired t-test, or a one-way ANOVA using Dunnett’s multiple comparison test (\**p* ≤ 0.05, \*\**p*≤ 0.01, *** *p*≤ 0.001, **** *p*≤ 0.001).

To determine whether the observed bacterial growth inhibition was due to prophage induction or due to direct antibacterial effects of the compounds, we quantified virus-like particle (VLP) production in all conditions where the decrease in bacterial growth was 15% ≤ AUC ≤ 75 %. In total, out of the 28/68 (41.17%) conditions that met this criteria, 10/28 (35.71%) led to significant increases in VLP production compared to the vehicle control (Fig. 1B), indicative of prophage induction^34^. Out of the 8 compounds tested, prophage induction was observed in at least one strain in at least 1 concentration for 5 compounds: taurocholic acid, hydrogen peroxide, peroxynitrite, human beta-defensin 3 (hBD3) and LEA (Fig. 1B). Hydrogen peroxide and peroxynitrite caused induction in 5/10 (50%) conditions and generally caused the greatest increase in VLP production (Fig. 1B). Interestingly, *B. caecimuris*, which contained only 1 predicted prophage, was not induced by any of the conditions tested (Fig. 1B). *Bacteroides* spp. generally have high sensitivity to bile salts^50^ and oxidative stress^51,52^,which could explain the lack of prophage induction observed in this strain (Fig. 1B). Both *A. muciniphila* strains were induced by hydrogen peroxide (Fig. 1B). However, the BAA-835 strain was uniquely induced by taurocholic acid and the YL44 strain was uniquely induced by peroxynitrite and LEA (Fig. 1B), suggesting that these two *A. muciniphila* strains have different sensitivity to prophage induction. *F. plautii* had broad sensitivity to the compounds tested, as it was induced by hydrogen peroxide, peroxynitrite, hBD-3 and taurocholic acid (Fig. 1B). Overall, significant increases in VLP production were observed in 10 conditions, across 3 bacterial strains and 5 distinct compounds, indicating that compounds characteristic of intestinal inflammation can trigger prophage induction in commensal gut bacteria.

An important consideration when quantifying VLPs using epifluorescence microscopy is that the presence of DNA contained within membrane vesicles or gene-transfer agents in cultures could influence reported viral concentrations^53^. We therefore verified increases in VLP production by extracting DNA from these *in vitro* cultures and measured the prophage:host ratio using the detected prophage regions shown in Supplemental Table S1. We proceeded with this approach in two representative conditions where we observed VLP increases using microscopy: *A. muciniphila* YL44 grown with 0.25 mM hydrogen peroxide and *F. plautii* YL31 grown with 2.5% taurocholic acid. For *A. muciniphila*, we observed a significant increase in the prophage:host ratios in *A. muciniphila* prophage Chantilly (Supplemental Fig. S1A). However, no significant increases in these ratios were observed for the two other characterized *A. muciniphila* prophages, Moulinsart and Chambord (Supplemental Table. S1), suggesting that induction is prophage specific in response to 0.25 mM hydrogen peroxide. Similarly, when *F. plautii* was grown with taurocholic acid, we observed an increase in prophage:host ratios with *F. plautii* phage Cormatin and a VIBRANT-predicted prophage (“prophage 1”, Supplemental Table. S1), but not the 4 other identified prophages (Supplemental Fig. S1B-C). It should be noted that despite observing significant increases in prophage:host ratios for specific prophages in *A muciniphila* and *F. plautii*, the effect sizes in both conditions were generally smaller than expected, suggesting that only a fraction of bacteria in the population are being induced.

### Dextran sodium sulfate (DSS)-colitis alters the composition of the OMM^12^ bacterial community

Knowing that inflammatory compounds can induce gut bacterial prophages *in vitro,* we next wanted to determine whether prophage induction could be detected at a community level *in vivo*. We used the OMM^12^ synthetic bacterial consortium^55,56^ to monitor changes in the mouse gut microbiota during intestinal inflammation. Following oral gavage of members of the OMM^12^ community to 15 adult germ-free (GF) mice, we administered 2% dextran sodium sulfate (DSS) for 5 days in the drinking water of 7 mice (co-housed in 3 cages) to induce colitis (Fig. 2A).

**Fig. 2.**
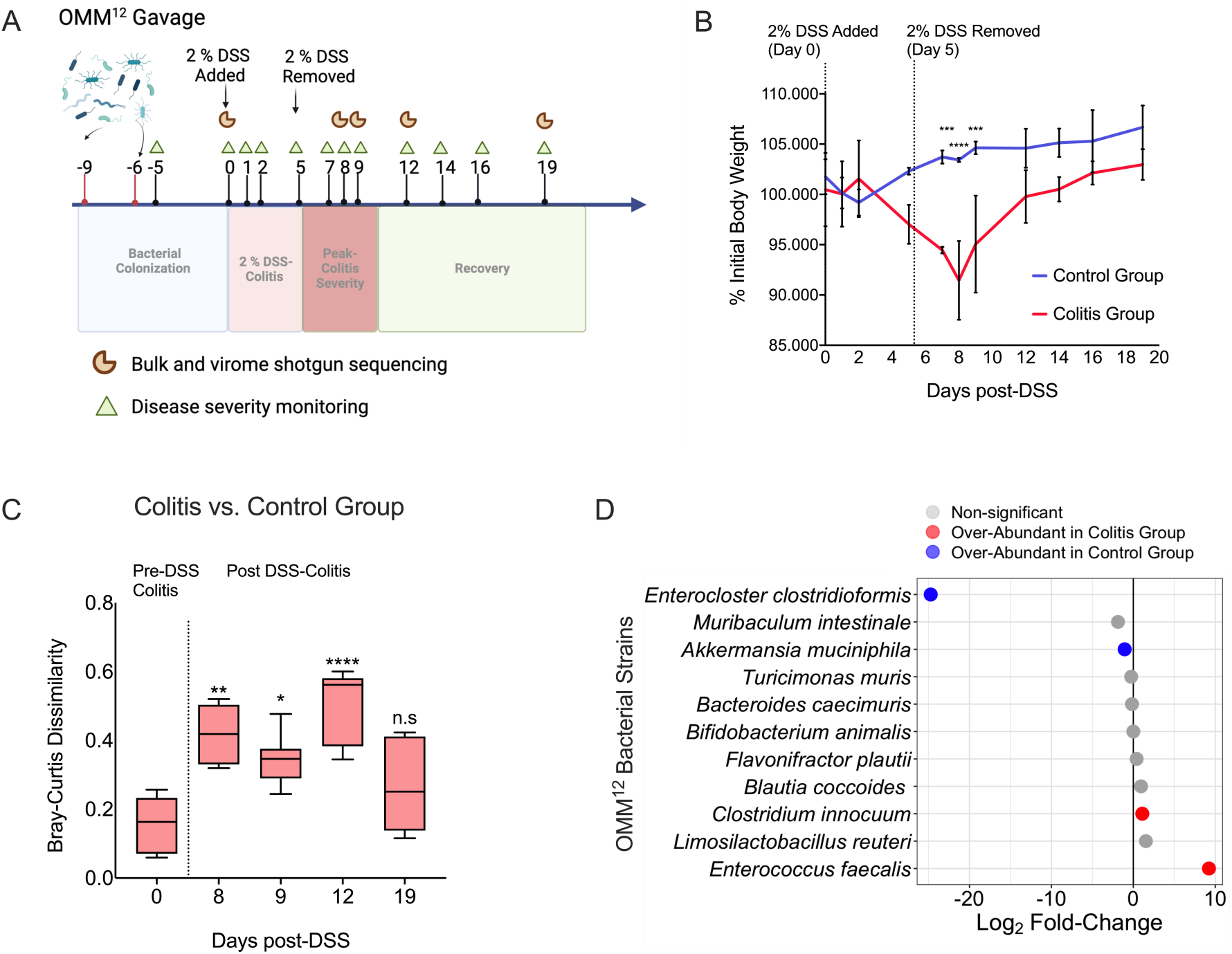
DSS-colitis shifts the composition of the OMM^12^ bacterial community. A) Schematic overview of the experimental study design. Fifteen mice were colonized with all OMM^12^ strains by oral gavage. The colitis group comprised of 7 female mice cohoused in 3 cages. The control group consisted of 6 male mice cohoused in 2 cages and 2 female mice cohoused in 1 cage. Two percent DSS was administered in the drinking water of 7 mice cohoused in 3 cages for 5 days (colitis group). Eight control mice cohoused in 3 cages remained on normal drinking water over the same period of time. Fecal samples within cages were collected and pooled (n=3 per treatment group) to extract viral and bulk DNA. B) Mean (± SD) percent initial body weight of mice was calculated in reference to weight 5 days pre-DSS. Significance was assessed between treatment groups using a repeated measures two-way ANOVA and Sidak’s multiple comparison test (*** *p*≤ 0.001, **** *p*≤ 0.0001). C) Bray-Curtis (BC) dissimilarity of the fecal bacterial community in mice between control and colitis groups was calculated from bulk metagenomes. Significance was assessed relative to baseline (day 0) using the Friedman test (\**p*≤ 0.05, \*\**p*≤ 0.01, **** *p*≤ 0.0001). D) The log_2_ fold-change of bacterial taxa was calculated between the colitis and control group during peak colitis (days 8 and 9 post-DSS). Taxa with a log_2_ fold-change ≥ 1 or ≤ −1 and an adjusted *p*-value ≤ 0.05 were considered differentially abundant using DESeq2 and the Wald test.

DSS-mediated colitis is reproducible and inducible, allowing us to follow changes in the microbiota over the course of inflammation. Further, compounds such as ROS^57^ and bile salts, associated with human IBD, are increased during DSS-colitis in mice, making it a suitable model to study prophage induction *in vivo.* Eight control mice (co-housed in 3 cages) not receiving DSS were also monitored across the same period and frequency (Fig. 2A). Fecal samples pooled per cage were collected throughout the experiment to monitor disease severity and track gut microbiota composition over time (Fig. 2A). The disease activity index (DAI) of mice peaked between days 5-9 post-DSS administration (Fig. S2A). Mice that received 2% DSS also experienced significant weight loss on days 7,8, and 9 post-DSS compared to control mice (Fig. 2B), suggesting heightened disease severity on these days. Although DAI decreased after day 9 post-DSS (Fig. S2A), mice exposed to DSS still had lower weights relative to control mice until the experimental endpoint at 19 days post-DSS (Fig. 2B).

To characterize phage-host dynamics in during colitis, we performed bulk metagenome and virome shotgun sequencing. However, we could not generate either bulk or virome DNA libraries during the DSS administration period and up two days after DSS removal (days 1-7 post-DSS), despite testing several cleaning protocols. This is likely due to DSS-mediated inhibition of DNA polymerase^58^, resulting in insufficiently low concentrations of Illumina DNA libraries (see methods). However, this did not prevent us from capturing the bacterial and viral communities when mice experienced peak colitis, i.e., days 8-9 post-DSS administration (Fig. 2B). Using the remaining available bulk metagenomes, we first characterized changes in bacterial community composition by calculating the Bray-Curtis (BC) dissimilarity between control and colitic mice over time. There was a significant increase in BC dissimilarity between these groups from baseline (day 0) to day 8 post-DSS (Fig. 2C, Fig. S2B), indicating that inflammation alters the composition of the OMM^12^ bacterial community. Significantly increased BC dissimilarity between control and colitic mice remained on days 9 and 12, and only partially returned to baseline levels by 19 days post-DSS (Fig. 2C, Fig. S2B), suggesting that these alterations are potentially related to disease severity. We next used DESeq2 to identify members of the bacterial community that were differentially abundant between colitic and control mice during peak colitis. Overall, there were strain-specific responses to inflammation (Fig. 2D, Fig. S2C-S2D): *Enterocloster clostridioformis* and *A. muciniphila* were significantly depleted during peak colitis, while *Enterococcus faecalis* and *Clostridium innocuum* were enriched (Fig. 2D, Fig. S2C-S2D). Of these bacteria, *E. clostridioformis* exhibited the most drastic response to inflammation, decreasing from a mean baseline relative abundance of 35.02% to close to, or below, the limit of detection in all colitic mice on days 8 and 9 post-DSS (Fig. 2D, Fig. S2C-S2D). The depletion of *E. clostridioformis* is in agreement with previous observations that some members of the *Lachnospiraceae* family exhibit increased sensitivity to DSS-inflammation^60,61^. In addition to the direct effects of inflammation on specific members of the community, complex inter-bacterial interactions^16,59^ are likely governing community composition of the OMM^12^ bacteria in response to inflammation.

### Dextran sodium sulfate (DSS)-colitis shifts the composition of the extracellular phageome in OMM^12^ - colonized mice

As we observed distinct changes to the composition of the bacterial community during DSS-colitis, we hypothesized that there would be similar changes to the phageome, due to prophage induction. Thirteen inducible prophages in the OMM^12^ community were recently described by Lamy-Besnier^43^. Since OMM^12^-colonized mice are free of lytic phages, any extracellular phages should be a consequence of prophage induction^43^. Using a shotgun viral metagenomics approach (see methods), we tracked the composition of these phages over the course of inflammation. At baseline, the extracellular phage community was predominantly composed of *Blautia coccoides* phage Montmirail, *B. caecimuris* phage Versailles, and *E. clostridioformis* phage Villandry (Fig. 3A). The prominence of these phages during baseline reflects the high relative abundances of their hosts and is likely an indication of basal levels of prophage induction for these specific phages. In some baseline and control samples, *B. caecimuris* phage Versailles in particular had high relative abundances, reaching up to 93.7 % of the community (Fig. 3A). However, this high relative abundance of phage Versailles was not observed across all baseline and control samples (Fig. 3A), suggesting that there is variance in the level of prophage induction, even in baseline conditions.

**Fig. 3.**
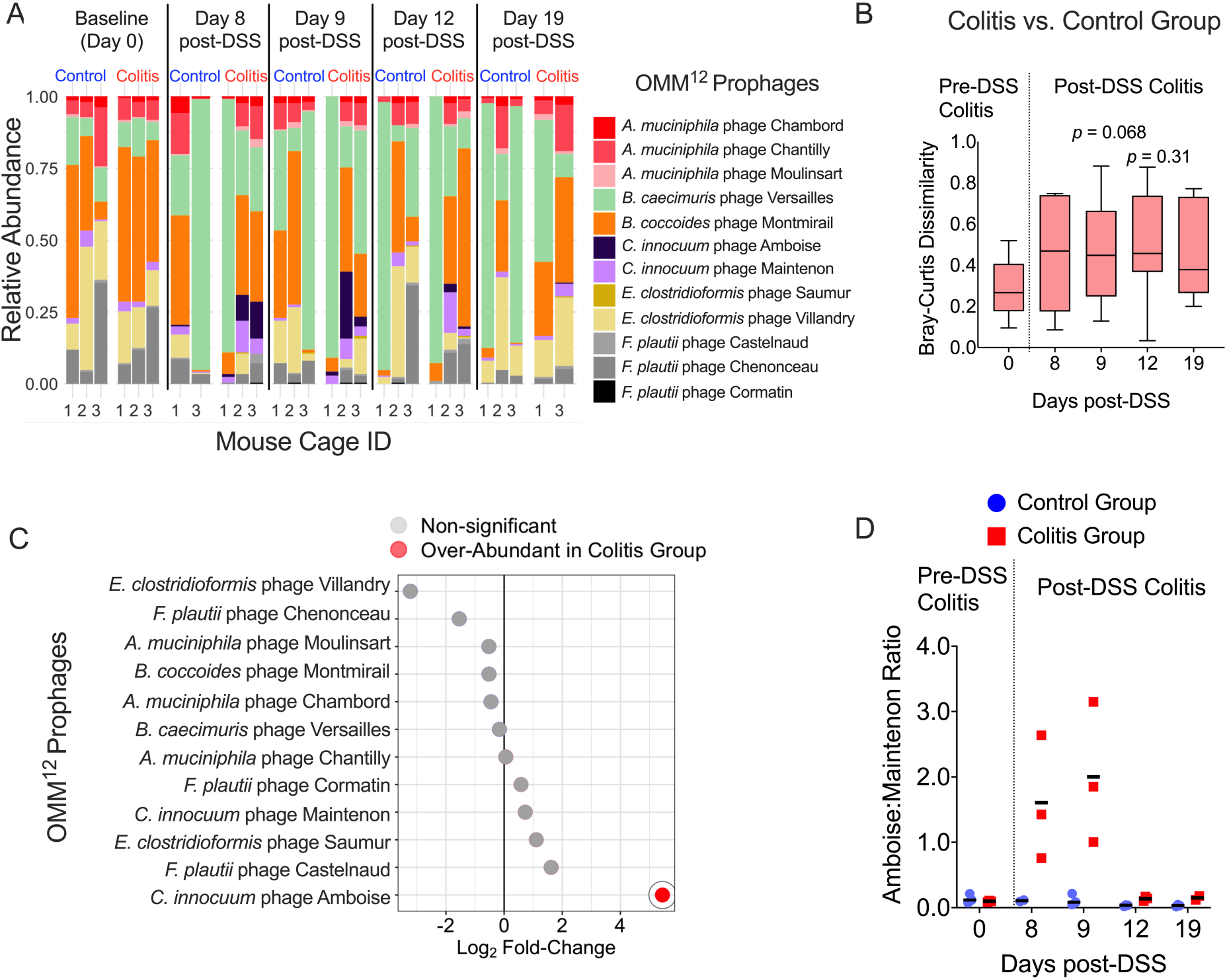
DSS-colitis shifts the composition of the extracellular phageome in OMM^12^ - colonized mice. Virome DNA was obtained from pooled fecal samples from the same cage (n=3 per treatment group). A) Relative abundance of free (extracellular) prophages within the OMM^12^ consortium. All prophages identified by Lamy-Besnier^43^ were included in analyses, except *Acutalibacter. muris* phage Fontainebleau since its host was not included in the initial GF mouse gavage. B) Bray-Curtis (BC) dissimilarity of the free phageome was calculated over time. Significance was assessed using the Friedman test (*p* ≤ 0.05). C) The log_2_ fold-change of free prophages in the OMM^12^ consortium was calculated between the colitis and control group during peak colitis (days 8-9 post-DSS). Phages with a log_2_ fold-change ≥ 1 or ≤ −1 and an adjusted *p*-value ≤ 0.05 were considered differentially abundant using DESeq2 and the Wald test. D) The length-normalized coverage ratios of *C. innocuum* phage Amboise: *C. innocuum* phage Maintenon. Horizontal lines represent group means. Significance was assessed between treatment groups using a repeated measures two-way ANOVA and Sidak’s multiple comparison test, but no significant differences were observed. Due to low biomass, viromes for 1 control sample (day 8 post-DSS) and 1 colitis group sample (19 days post-DSS) could not be obtained. For these days significance could not be assessed (panels B and D).

Despite this variance, we detected a modest increase in phageome BC dissimilarity between control and colitic mice from baseline to peak colitis (9 days post-DSS, *p* = 0.068) (Fig. 3B, Fig. S3A), indicating that the temperate phageome is altered in response to DSS-colitis. Due to low fecal biomass for some cages (control mouse cage “2” and colitis mouse cage “2”), virome samples were missing on days 8 and 19 post-DSS, and therefore significance could not be assessed for these days. Still, there was a trending increase in dissimilarity on days 12 and 19-post DSS (Fig. 3B, Fig. S3A), suggesting that alterations to the temperate phageome remain after peak colitis.

In terms of specific phages, there was a drastic increase in *C. innocuum* phage Amboise during peak colitis (Fig. 3C, Fig. S3C, log_2_ fold-change increase = 5.42), which was deemed differentially abundant using DESeq2 (Fig. 3C, Fig. S3B). The relative abundance of this phage peaked on days 8 and 9 post-DSS but remained higher in colitic mice at all post-DSS time points (Fig. 3A, Fig. S3C). Based on these data, phage Amboise may expand during inflammation as a consequence of increased prophage induction. Alternatively, since its host also expanded during inflammation (Fig. 2D), the increase in phage Amboise may simply by a consequence of the enrichment of its host. To distinguish between these two scenarios, we determined the ratios of *C. innocuum* phage Amboise to *C. innocuum* phage Maintenon. If the expansion of phage Amboise was simply due to expansion of *C. innocuum,* the ratios of phage Amboise to Maintenon should remain similar over the course of inflammation. Instead, there was an increase in the ratio of phage Amboise: phage Maintenon during peak colitis (Fig. 3D), suggestive of increased induction of phage Amboise. While not statistically significant, all mice in the colitis group exhibited a > 10-fold increase in these ratios from baseline to peak colitis (Fig. 3D). We also used PropagAtE^62^ to determine the prophage:host ratios of phage Amboise from bulk metagenomes as an indicator of prophage activity. There was a modest, but non-significant increase in the prophage:host ratio of phage Amboise in the colitis group during peak colitis (Fig. S3D). Interestingly, the prophage:host ratio of phage Amboise remained below 1 throughout the study (Fig. S3D), indicating that a subset of the *C. innocuum* population may not contain the integrated Amboise prophage. During inflammation, it is possible that selection against prophage-containing members of the bacterial population is taking place to limit deleterious consequences of strong induction^63^. This selection could therefore limit the use of prophage:host ratios as an estimate for prophage activity^62^, and could potentially explain why large increases in this ratio for phage Amboise were not observed, despite its expansion in the extracellular phageome. Due to insufficient sequence coverage, we could not definitively determine the proportion of *C. innocuum* variants lacking the integrated phage Amboise during peak colitis. Lastly, to validate that any prophage induction in *C. innocuum* was not due to direct bacteria-DSS interactions, we grew *C. innocuum in vitro* with 2% DSS and confirmed that DSS did not increase VLP production compared to the vehicle control (Fig. S3E).

While phage Amboise was the only differentially abundant phage between colitic and control mice during peak colitis (Fig. 3C, Fig. S3B), we also report a non-significant reduction in the *E. clostridioformis* phage Villandry (Fig. 3C, log_2_ fold-change = −3.22), likely as a consequence of the depletion of its host (Fig. 2D). Interestingly, despite the depletion of *E. clostridioformis* below the limit of detection, phage Villandry was still detected during peak colitis in the colitis groups in 2/3 cages at relative abundances up to 12.23 % (Fig. S3F).

Similarly, there was increased relative abundance of *E. clostridioformis* phage Saumur during peak colitis in 2/3 cages in the colitis group (Fig. S3G). Given the low coverage of *E. clostridioformis,* it is difficult to determine definitively whether the presence of phage Villandry and phage Saumur during peak colitis is an indication of heightened prophage induction during inflammation, or simply the fact that these phages can persist at low levels even in the absence of detection of their hosts. Regardless, our data collectively suggest increased prophage induction in certain bacterial taxa in response to DSS-colitis.

### Murine T-cell colitis shifts the composition of the extracellular temperate phageome

We next investigated whether changes in the extracellular temperate phageome could be detected during inflammation in a T-cell model of colitis, in part to validate our previous observations in DSS-colitis and to study these changes in conditions more clinically and immunologically relevant to IBDs^64^. We thus reanalyzed a previously published viral metagenome dataset^65^, where 3 C57BL6/J *Rag1^−/−^* mice with a conventional laboratory microbiota experiencing T-cell colitis and 3 controls were sampled at 3 time points over 6 weeks (Fig. S4A).Mice experienced peak colitis severity at day 42 (Fig. S4A)^65^. We applied a viral metagenomics approach to this dataset to assemble extracellular phage scaffolds and assign putative replication cycles (see methods). Considering dereplicated scaffolds with > 50 % completeness as viral operational taxonomic units (vOTUs)^66^, we identified 81 of the 143 unique vOTUs identified across all 6 mice as temperate. We also assigned a bacterial host to each vOTU using iPHoP^67^. Since the inference of phage hosts is biologically informative, we considered predicted phage host family (PHF) to be a distinct taxonomic unit, of higher rank than a vOTU^68^. For example, we consider a *Lactobacillaceae* PHF to correspond to the group of vOTUs predicted by iPHoP to infect bacteria from the *Lactobacillaceae* family.

Of the 143 unique vOTUs, 109 were assigned a PHF. These 109 assignments represented a total of 22 unique PHFs (Fig. S4B). There were distinct differences in the PHFs belonging to the predicted temperate and virulent fractions of the phageome, with 59% (13/22) of PHFs found exclusively in the temperate fraction of the phageome (Fig. S4B). Only two PHFs, *Enterobacteriaceae* and *CAG-552*, were exclusively found in the virulent phageome (Fig. S4B). As observed previously^65^, using our metagenomics pipeline we also found that there was divergence in the phageome over the course of colitis, peaking at day 42 when mice were experiencing high disease severity (Fig. S4C).

Compared to control mice, there was a modest, non-significant increase in the relative abundance of predicted temperate phages during peak colitis (Fig. 4A). Despite this, the temperate subset of the phageome diverged over time in response to colitis, particularly at day 42, as determined by BC dissimilarity of the phageome at the vOTU level (Fig. 4B). While control and colitis groups also diverged in the virulent subset of the phageome, this occurred to a lesser degree (Fig. 4C). At day 42, the BC dissimilarity between control and colitis mice was significantly higher in the predicted temperate phageome compared to the virulent phageome (Fig. 4C). Similar trends were noted when comparing predicted temperate and virulent phageome divergence at the PHF level of taxonomy (Fig. S4D). These data suggest that changes to the composition of the whole free phageome during inflammation are largely driven by extracellular temperate phages.

**Fig. 4.**
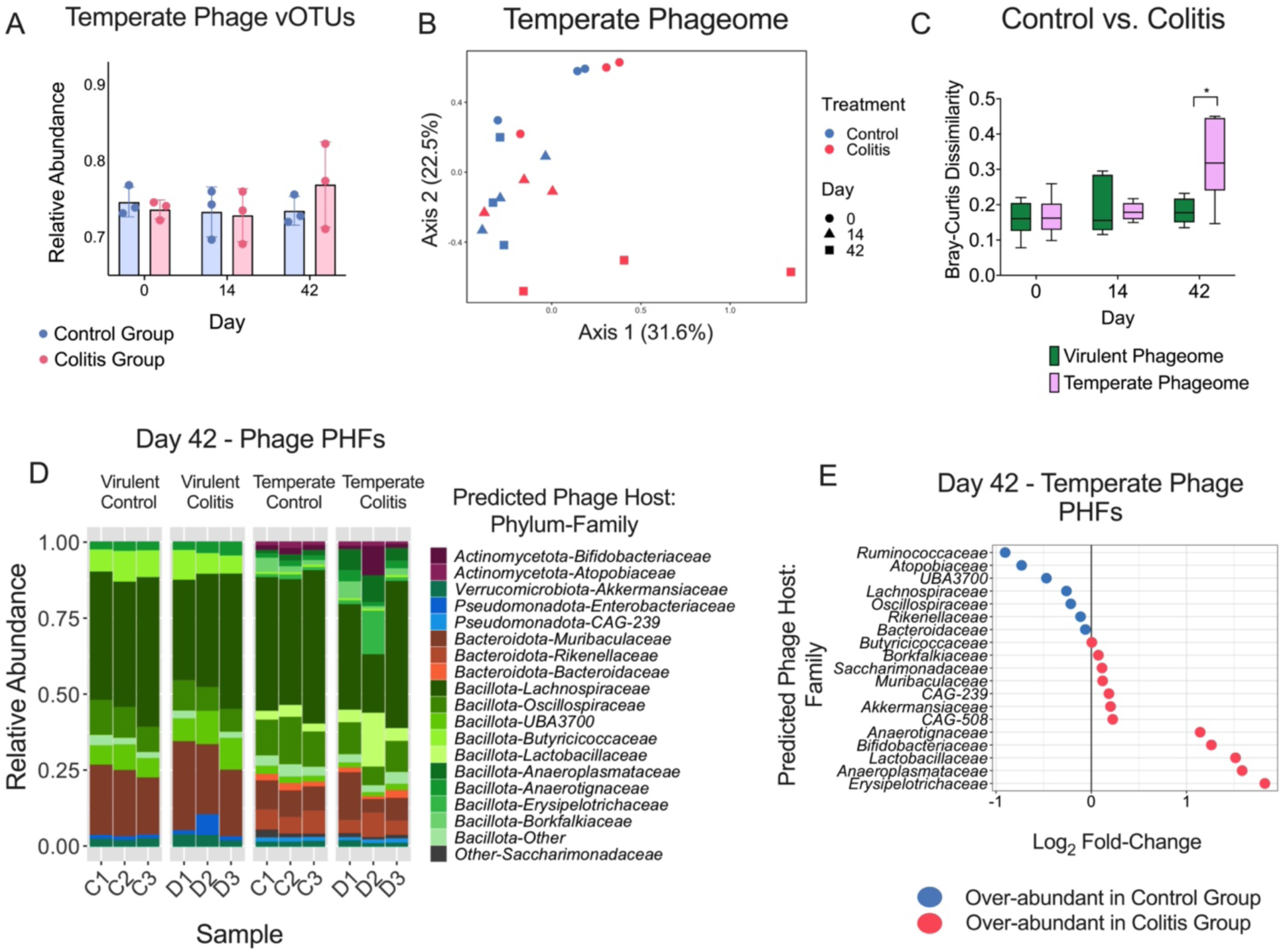
Murine T-cell colitis shifts the composition of extracellular temperate phages. Fecal viromes were obtained from publicly available data generated by Duerkop^65^ (n=3 mice per treatment group). A) Mean relative abundance (± SD) of temperate viral OTUs (vOTUs) in the control group and the T-cell colitis group. vOTUs were identified as temperate using bacphlip^69^. B) PCoA on Bray-Curtis (BC) dissimilarity at the phage vOTU level of taxonomy. C) BC dissimilarity between the control and colitis group in the temperate and virulent fractions of the free phageome at the vOTU level of phage taxonomy. Significance was assessed using a repeated measures two-way ANOVA using Sidak’s multiple comparison test (* *p* ≤ 0.05). D) Relative abundance of predicted phage host families (PHFs) on day 42 during peak colitis. “C” and “D” on the x-axis refers to the ID of mice from the control and disease (colitis) groups respectively. E) Log_2_ fold-change of temperate phage PHFs between the colitis group and control group on day 42 during peak colitis. iPHoP^67^ was used for bacterial host prediction. Note that over-abundance here does not correspond to statistical significance.

To determine which vOTUs were most altered inflammation, we performed DESeq2 analysis. Two vOTUs significantly increased during peak colitis in the T-cell colitis group compared to controls (Fig. S4E). Both vOTUs were bioinformatically identified as temperate: one with an unidentified host and another predicted to infect *Anaerotignaceae.* When considering PHFs, there was also divergence in the temperate fraction of the phageome during peak colitis (Fig. 4D). Specifically, there was an enrichment in 5 temperate PHFs during colitis: *Bifidobacteriaceae, Erysipelotrichaceae*, *Lactobacillaceae*, *Anaerotignaceae*, and *Anaeroplasmataceae* (Fig. 4D-E). Phages predicted to infect *Lactobacillaceae* and *Anaeroplasmataceae* were enriched across all three mice (Fig. 4D, Fig. S4F-G), while the *Anaerotignaceae* PHF was enriched in two mice (Fig. 4D, Fig. S4H) and *Erysipelotrichaceae* and *Bifidobacteriaceae* PHFs (Fig. 4D, Fig. S4I-J) were enriched in only one mouse. The phageome of mouse “D2”, previously shown to exhibit high variance during peak colitis^65^, also diverged to a large degree at the PHF level in our analyses of the temperate phageome, with high relative abundances of *Lactobacillaceae*, *Erysipelotrichaceae,* and *Bifidobacteriaceae*-infecting phages (Fig. 4D). Compared to the temperate phageome, there were minimal changes in the predicted virulent PHFs, with the exception of an expansion of *Enterobacteriaceae-*infecting virulent phages in mouse “D2” (Fig. 4D). Together, these data suggest that specific PHFs drive the shifts in the temperate phageome during murine T-cell colitis, possibly resulting from induction of their bacterial hosts in response to inflammation.

Given the evidence of prophage induction we observed in mouse models of intestinal inflammation, we next explored whether similar changes would be observed in a human IBD cohort. Previous studies have shown that temperate phages are generally enriched in IBD patients compared to non-IBD controls^39,41,70^. However, these studies have not determined the contribution of temperate and virulent phages to overall changes in phage diversity. We re-analyzed a previously published dataset from the human microbiome project 2 (HMP-2)^6^, containing longitudinal bulk metagenome samples from 65 CD patients, 38 UC patients, and 27 non-IBD controls. Using the same viral metagenomics approach used to analyze the murine T-cell colitis data set earlier, we employed a similar approach here. We first removed samples which contained low read counts. In total 1,093 samples from 57 CD, 31 UC, 27 non-IBD controls remained for further analyses. Across these individuals, we assembled 3,862 distinct vOTUs. Of these, 2,176 (56.34 %) were identified by bacphlip as temperate.

Splitting the dataset according to predicted replication cycle, we observed that disease diagnosis status generally explained a low proportion of virome variance in both the temperate (Fig. S5A; R^2^ = 0.022) and virulent subsets (Fig. S5B; R^2^ = 0.026) of the whole phageome. Lloyd-Price *et al.* defined dysbiotic samples within this HMP-2 dataset as those with high microbiota divergence from non-IBD controls^6^. Compared to diagnosis status, dysbiosis status did explain more virome variance in both the temperate (Fig. S5C; R^2^= 0.037) and virulent subsets (Fig. S5D; R^2^ = 0.029) of the whole phageome.

Despite the modest alterations in the virome at the whole community level, we next determined whether specific PHFs were differentially abundant based on dysbiosis status. To assess whether replication cycle drove differences in differential abundance, analyses was performed on both temperate and virulent subsets of each PHF (Fig. S5E). In total 4 distinct PHFs were differentially abundant: *Acutalibacteraceae, CAG-74*, *Ruminococcaceae*, and *Enterobacteriaceae* (Fig. S5E). For the *Ruminococcaceae*, and *Enterobacteriaceae* PHFs, both the virulent and temperate subsets were differentially abundant (Fig. S5E). Interestingly, for *Enterobacteriaceae* PHF, the temperate subset was more enriched compared to the virulent subset (Fig. S5E; temperate-to-virulent fold-change ratio = 3.38), which could be indicative of induction or increased prophage maintenance in this bacterial family. For the *Acutalibacteraceae* and *CAG-74* PHFs, only the temperate subsets were significantly depleted. These data together suggest that different phage replication cycle dynamics may explain the differential abundance of these phages. Thus, while the temperate and virulent subsets of the phage community, as a whole, shift to similar degrees in the context of IBD samples, the resulting dynamics of certain phage taxa may be dependent on specific phage-host interactions.

## Discussion

As phages are strong regulators of bacterial communities in the gut, alterations in their diversity are relevant to several diseases, including IBDs. However, the factors behind these changes and the role of phage replication strategies in driving such alterations remain poorly understood. Here, using a combination of *in vitro* prophage induction assays, a murine model of IBD, and re-analyses of previously published metagenomic datasets, we provide evidence that a switch from lysogenic replication to lytic replication could contribute to these changes in phage community composition during intestinal inflammation.

Extensive changes to the metabolome and the gut environment are associated with IBDs^6,12^. In agreement with Fornelos *et al.,*^48^ we showed that several compounds associated with the gut inflammatory response are directly inhibitory to commensal gut bacterial strains. Of the conditions that led to bacterial growth inhibition, we found that 35.71% led to increases in VLP production. The two *A. muciniphila* strains, BAA-835 and YL44, exhibited differences in sensitivity to prophage induction upon exposure to the same inflammatory compounds. These differences are indicative of strain-specific differences in sensitivity to stress, or possibly due to differences in phage-phage interactions, as has been observed previously in *E. faecalis* polylysogenic strains^71^.

Hydrogen peroxide and peroxynitrite were particularly strong prophage inducers of the bacterial isolates screened. Elevated levels of ROS are associated with IBDs^72^, likely as a consequence of increased recruitment of ROS-producing neutrophils^73,74^ and increased production of ROS by colonocytes^44^. High levels of RNS are also associated with intestinal inflammation^45,75^, due to upregulation of iNOS^76^. That these compounds are strong inducers is consistent with the fact that ROS and RNS are known to cause bacterial DNA damage, a potent trigger of prophage induction through activation of the SOS stress response^29^. Notably, Diard *et al.* demonstrated that prophage induction *in Salmonella Typhimurium* occurs in response to intestinal inflammation, and importantly ROS and RNS were shown to induce these Salmonella prophages *in vitro*^77^. A number of prophage induction triggers have been identified in the gut^28^, including host hormones, dietary compounds^36,37^, and several xenobiotics^28^. Our data build on these findings to suggest that the products of the intestinal inflammatory response can similarly trigger prophage induction. Still, future studies will be needed to test whether these specific inducing agents have similar effects at a community level *in vivo*. For instance, induction was not observed in *B. caecimuris* I48 *in vitro*, despite high activity of its prophage Versailles in OMM^12^-colonized mice. In addition to measuring VLP production by epifluorescence microscopy, we calculated prophage:host ratios in select conditions, and determined that in both*A. muciniphila* and *F. plautii*, increased ratios were observed for some, but not all prophages. Different co-residing temperate phages may be under different regulatory cues, and may have different burst sizes, replication rates, or packaging rates^78^, which could all explain why some replicate more successfully than others in response to these compounds.

At a community level, there was evidence of prophage induction in response to inflammation in two distinct mouse models of colitis. Most studies that have investigated the phageome in inflammation have cross-sectionally characterized changes in the composition of the community in health and in disease. Using a tractable model of inflammation, we could follow how the phageome changes longitudinally during disease progression. Using OMM^12^-colonized GF mice was ideal for studying prophage induction at a community level: these strains have complete genome assemblies available and contain experimentally validated prophages^42,43^, which allows for confident phage-host matching compared to assembly-based methods.

Importantly, since these OMM^12^-colonized mice are free of virulent and eukaryotic viruses^43^, we can also conclude that any changes in the virome in this system are a consequence of temperate phages released via prophage induction. Recent findings by Munch *et al*. suggest that antibiotic administration can trigger prophage induction in OMM^12^-colonized mice^59^, supporting the utility of this model for investigating how various perturbations can impact prophage induction in the gut.

While community-level alterations in the extracellular OMM^12^ phageome during DSS-colitis were less drastic than in the bacteriome, shifts were still observed in specific phage taxa. In particular, there was a striking increase in *C. innocuum* phage Amboise, likely resulting from increased prophage induction. Lamy-Besnier reported low levels of Amboise prophage activity in OMM^12^-colonized mice, but high prophage activity *in vitro*^43^. These observations suggest that the activity of this prophage is highly contextual and linked to changes in the gut environment during inflammation. In agreement with the changes in the extracellular temperate phageome in response to DSS-colitis, we also observed distinct changes in the composition of the predicted temperate phageome during the progression of murine T-cell colitis. Importantly, the temperate fraction of the free phageome shifted to a larger degree than the virulent fraction between the control and colitis groups, suggesting that the alterations in phage diversity in response to T-cell colitis are largely due to prophage induction and subsequent release of extracellular temperate phages. The fact that changes to the gut phageome were observed during inflammation in two mechanistically distinct models of colitis provides strong evidence that inflammation may broadly disrupt the longitudinal stability characteristic of the gut phageome^33,79^. During homeostasis, passive replication of integrated prophages, low levels of spontaneous prophage induction^33^, and replication by phages that exhibit “carrier-state” replication^80^ dominate the gut phageome and likely contribute to community stability^33^. These dynamics are consistent with the Piggyback-the-Winner (PtW) model, where lysogeny is favoured at high bacterial densities^32^.

From an ecological perspective, our data suggest that there may be a shift from lysogeny and PtW dynamics to lytic replication over the course of inflammation. Importantly, in both colitis models, alterations in the gut phageome coincided with peak disease severity, suggesting that that these shifts to lytic replication occur in tandem with changes to the gut microenvironment during inflammation.

In agreement with our observation that the temperate phageome shifts in response to inflammation in mice, several studies have reported an enrichment in extracellular temperate phages in CD and UC patients compared to non-IBD controls^39–41,70^. While analyses of the HMP-2 dataset confirmed that there were minor shifts to the temperate phageome in IBD and dysbiotic samples, we noted that the shifts in the virulent and temperate phageomes were similar. As the samples in the HMP-2 cohort were obtained at various time intervals over the course of a year, it is possible that metagenomic evidence of prophage induction may be transient and therefore missed by extended sampling intervals. In support of this idea, Sutcliffe *et. al* showed that over the course of 2.4 years, triggered prophage induction events in a healthy individual are rare^33^.

While these events are likely more frequent in IBD patients, they may still be difficult to detect with infrequent sampling. The dysbiotic samples identified by Lloyd-Price reflect periods of heightened microbiota dissimilarity^6^, but even this designation seems insufficient to detect subtle differences in prophage activity. As mouse models of inflammation tend to exaggerate disease severity compared to human IBDs^81^, this disparity may also explain the heightened prophage induction we observed in these experimental models. Future studies investigating gut microbiota changes in IBDs should thus employ sampling strategies with short intervals, ideally centered around periods of high disease severity when possible. Despite these limitations, using the extracellular phageome, we observed that differential abundances of some PHFs were driven by either virulent or temperate phages. For instance, the enrichment in *Enterobacteriaceae*-infecting phages was largely driven by temperate phages, indicating that prophage induction could be responsible for these dynamics. As *Enterobacteriaceae* increase in response to inflammation^6^, the release of temperate phages may be a means to promote lysogenic conversion and the spread of virulence factors, allowing these bacteria to adapt to the inflamed gut and cause disease^77^. Still these data should be interpreted with caution, as using a bulk metagenomics approach we could not distinguish between integrated and excised prophages.

Taken together, our data suggest that at a community level, prophage induction may occur to a higher degree in certain bacterial taxa compared to others. Interestingly, despite high levels of prophage activity, these bacterial taxa exhibit diverse patterns of abundance. For instance, the expansion in temperate *Enterobacteriaceae-*infecting phages in the HMP-2 dataset and the expansion of phage Amboise in DSS-colitis were associated with an expansion of their hosts. In contrast, the enrichment of *Anaerotignaceae*-infecting phages in T-cell colitis coincided with a decrease in host abundance (data not shown), and we also noted the presence of *E. clostridioformis* phages during peak DSS-colitis despite the depletion of their host. These observations are in line with data describing the enrichment of Bacillota-infecting temperate phages in IBD patients^39,40^. Cornuault also described that *Faecalibacterium prausnitzii*-infecting temperate phages are particularly abundant in IBD microbiomes, despite the depletion of their hosts^40^. Therefore, the shifts in temperate phage composition across different host abundance dynamics reflects complex ecological scenarios that are likely specific to certain phage-host pairs.

While our findings shed important light on phage replication cycle dynamics, they may also be relevant to disease progression. Indeed, previous work demonstrated that extracellular phages from UC patients exacerbate intestinal inflammation in mice^22^. The release of free temperate phages as a result of prophage induction could therefore contribute to heightened pro-inflammatory responses and should therefore be further investigated. Overall, by combining *in vitro* and whole community bioinformatics approaches, we link changes in the gut environment during inflammation to prophage induction and alterations in the phageome. Clarifying the mechanisms that underly alterations in the phageome is crucial in understanding how phages regulate bacterial communities during disease and consequently impact human health.

## Resource Availability

### Lead contact

### Materials availability

This study did not generate unique reagents

### Data and code availability

Virome and bulk metagenome reads are available at XXX and are publicly available as of the date of publication. All original code has been deposited at https://github.com/anshulsinha1/IBD_induction_manuscript and is publicly available as of the date of publication.

## Author Contributions

A.S designed and performed experiments, analyzed results, and wrote the manuscript. A.Q assisted in growth curve and VLP quantification. T.B provided technical assistance for DNA extraction and library preparations. A.S and C.F.M conceived the project, designed experiments, edited, and wrote the manuscript. The authors read and approved the final manuscript.

## Acknowledgments

We thank all members of the Maurice lab for their continuous feedback and criticism on all aspects of this work. A great thanks to the McGill Centre for Microbiome Research for their assistance in carrying out the OMM^12^ mouse experiments. We would especially like to thank Cynthia Faubert and Catherine Hudson for their technical expertise and support. Microscopy was performed using the McGill University Advanced Bioimaging Facility (ABIF).

## Ethics approval

All mouse experiments were carried out in accordance with the approved McGill University animal use protocol (7999).

## Supplemental information

Document S1. Figures S1-S5 and Table S1

## STAR Methods

### Experimental model and study participant details

#### Model of 2% dextran sodium sulfate (DSS)-colitis in OMM^12^-colonized mice

Fifteen GF C57BL/6 mice 6-8 weeks in age were purchased from Charles River and maintained in flexible film isolators at the McGill Centre for Microbiome Research animal facility. In total, the colitis group comprised of 7 female mice cohoused in 3 cages. The control group consisted of 6 male mice cohoused in 2 cages and 2 female mice cohoused in 1 cage. All mice had unlimited access to autoclaved mouse breeder’s diet and water. Mouse experiments were carried out in accordance with the approved McGill University Animal Use Protocol #7999.

A schematic of the experimental timeline and sampling schedule is shown in Fig. 2A. To colonize GF mice, the 11/12 bacterial strains belonging to the OMM^12^ consortium were prepared at equal concentrations and combined, according to Eberl^55^. Notably, *Acutalibacter muris* was excluded from the consortium in this model due to insufficient *in vitro* growth during preparation. Briefly, the remaining strains were grown anaerobically (Mandel Scientific Company Inc., Guelph, ON, Canada; 5 % Hydrogen, 20 % Carbon Dioxide, 95 % Nitrogen) at 37°C for 1-2 days in anaerobic *Akkermansia* media (AAM) or brain infusion broth (BHI) media^34^ (Thermo-Scientific) supplemented with hemin (5 μg/mL), vitamin K (1 μg/mL) and 0.05 % L-cysteine. Following incubation, the OD_600_ across all strains was normalized and equal volumes of each strain were added to a cryopreservation tube containing glycerol (10% final concentration). OMM^12^ preparations were thawed and administered to each mouse by oral gavage (150 uL). The same oral gavage was repeated 72 hours later to improve colonization stability^55^. Nine days following the first OMM^12^ gavage, 2 % DSS (MP biomedicals) was administered in the drinking water of the 3 cages in the colitis group for 5 days, after which the mice returned to drinking water without DSS. Following DSS removal, mice were monitored for an additional 14 days. Control mice received drinking water without DSS throughout the time course. Fecal samples were collected at various time points (Fig. 2A) and stored at −80 °C prior to DNA extraction.

### Monitoring dextran sodium sulfate (DSS)-colitis severity in OMM^12^ - colonized mice

Weight loss, stool consistency and the presence of blood in stool were assessed to monitor the severity of DSS-colitis. The percent weight loss of each individual mouse was measured over the course of the experiment and measured in reference to weight 5 days prior to DSS administration. A weight loss score^82^ was assigned to each mouse (0 = no weight loss, 1 = 1-10 % weight loss, 2 = 10-15 % weight loss, 3 = 15-20 %, 4 = > 20 %). The consistency of pooled fecal samples from each cage was measured blinded on a scale from 0-4: (0 = dry stool, 2 = loose stool, 4 = diarrhea)^82^. The presence of blood in pooled fecal samples was determined blinded using the Hemoccult SENSA kit (Beckman Coulter) and measured on a scale from 0-4 (0 = absence of blood, 2 = positive hemoccult test, 4 = positive test with visible evidence of rectal bleeding)^82,83^. The DAI was calculated as the sum of weight loss, stool consistency and hemoccult scores^82^.

## Method details

### Growth conditions for in vitro prophage induction assays

Four bacterial strains (Table. S1) were grown anaerobically (Mandel Scientific Company Inc., Guelph, ON, Canada; 5 % Hydrogen, 20 % Carbon Dioxide, 95 % Nitrogen). *A. muciniphila* YL44, *A. muciniphila* BAA-835 and *B. caecimuris* I48 were grown in BHI supplemented with hemin (5 μg/mL), vitamin K (1 μg/mL) and 0.05 % L-cysteine. *F. plautii* YL31 was grown in AAM. Strains were grown overnight at 37°C and then sub-cultured to a starting OD_600_ = 0.1 (n=3). Subcultures were grown in 96-well plates with inflammatory compounds of interest (Table. 1) or the corresponding vehicle controls to reach a final volume of 200 uL. Bacterial growth was monitored by measuring the OD_600_ (Epoch 2 microplate spectrophotometer, Biotek Instruments) until stationary phase. The AUC for each condition compared to its vehicle control was calculated using GraphPad Prism (v.8.4.3).

### Virus-like particle (VLP) enumeration for in vitro prophage induction assays

Conditions where the AUC decrease was ≥ 15 % and ≤ 75 % were selected for VLP enumeration. For these conditions, samples at stationary phase were fixed with 1% formaldehyde for 15 min. Samples were then diluted (between 1:10 – 1:100) in Tris-EDTA (TE) buffer and filtered to remove bacterial cells, using sterile 0.2 μm syringe filters (Millex-GP, Millipore Sigma). For each biological replicate, VLPs were filtered in duplicate onto 0.02 μm filter membranes (Anodisc, GE Healthcare), and stained with SYBR-Gold (2.5X concentration for 15 min). For each VLP filter, a minimum 300 VLPs, or 25 fields of view, were counted using an epifluorescence microscope at a 1,000X magnification (Zeiss, Axioskop).

### Prophage:host ratio-based validation of in vitro prophage induction assays

To validate increases in VLP production observed by epifluorescence microscopy, prophage:host ratios were determined in two conditions; 1) *A. muciniphila* grown with 0.25 mM hydrogen peroxide and 2) *F. plauti* grown with 2.5% taurocholic acid. Cultures were grown anaerobically at 37°C in triplicate until stationary phase. Five hundred microlitres of culture supernatant were added to 500 uL of inhibitEX buffer (Qiagen). Bulk DNA was extracted using the QIAamp Fast DNA Stool Mini kit (Qiagen), according to the manufacturer’s instructions.

### Bulk DNA extractions from OMM^12^-colonized mice

Bulk DNA was extracted from pooled feces from each cage, as described previously^79^. Briefly, stool was thawed, resuspended in 1 mL of inhibitEX buffer (Qiagen), and transferred to 2 mL tubes containing 0.5 g of 0.1 mm zirconium beads (Biospec), 0.7 g of 1 mm zirconium beads (Biospec), and 1 3.5 mm glass bead (Biospec). Samples were homogenized by bead-beating (Biospec) for 30 sec twice, allowing to cool on ice for 1 min in between each homogenization. Bulk DNA was then extracted from the resulting fecal suspension, using the QIAamp Fast DNA Stool Mini kit (Qiagen), according to the manufacturer’s instructions.

### Virome DNA extractions from OMM^12^-colonized mice

VLP DNA was extracted from fecal samples pooled within each mouse cage. Pooled fecal pellets were thawed, resuspended in 1mL sterile PBS and thoroughly vortexed until homogenized. The fecal suspension was centrifuged at 1,000*g* for 1 min at room temperature. The resulting supernatant was centrifuged at 10,000*g* for 15 min at room temperature to separate bacterial and viral communities. The VLP-containing supernatant was then centrifuged in 0.2 μm UltraFree-MC centrifugal filters at 12,000*g* for 2 min at room temperature to further remove bacterial cells. Chloroform (18% final concentration) was added directly to the extracted VLPs, and the solution was thoroughly vortexed and centrifuged at 21,000*g* for 5 min. The upper layer was removed, treated with Turbo DNAse (10 U) and Benzonase (125 U) and incubated for 1.5 hrs at 37 °C. To inactivate the DNAse cocktail, 19 mM EDTA was added and the solution was incubated for 30 min at 75°C. DNA was then extracted from the resulting VLPs using the QIAamp Mini Elute Virus Spin Kit (Qiagen), doubling the volume of buffer AL, protease, and ethanol indicated in the manufacturer’s instructions.

### Sequencing of bulk and VLP DNA from OMM^12^-colonized mice

Shotgun sequencing libraries from extracted bulk and VLP DNA were prepared using the Illumina DNA prep kit, according to manufacturer’s instructions. Notably, bulk metagenome and virome libraries could not be prepared from days 1-7 post-DSS administration. During this period, DSS likely is in sufficient concentrations in mouse fecal pellets to inhibit DNA polymerase, as has been previously described^58^. Libraries could also not be prepared for 1 control sample (day 8 post-DSS) and 1 colitis group sample (19 days post-DSS) due to low fecal biomass. The concentration of the remaining libraries were assessed using the Qubit HS kit (Life Technologies). Libraries were normalized, pooled, and sequenced using a Novaseq X Plus, generating 150 bp paired-end reads. On average, 6.3 million paired-end phageome reads and 13.6 million paired-end bulk metagenome reads were generated per sample.

### Characterization of bacterial communities in the OMM^12^-colonized mice

Raw reads from bulk metagenomes were trimmed based on sequence quality, and adapter sequences were removed using Trimmomatic (v 0.33)^84^. Quality-controlled reads that aligned to mouse genomes were removed by Bowtie2 (v 2.2)^85^. MetaPhlAn (v. 4.0, db version vOct22) was used on the remaining reads to generate counts tables and obtain relative abundances. MetaPhlAn^86^ was used in order to avoid spurious mapping of reads to reference assemblies and to confirm that only bacteria in the OMM^12^ consortium were present in fecal samples. Assemblies of each bacterial strains in the OMM^12^ consortium were obtained from Garzetti^87^.

### Characterization of phage communities in the OMM12-colonized mice

Raw reads from the extracellular phageome were quality-controlled as described above. Prophage coordinates and the genome sequences of 13 previously identified prophages in the OMM^12^ consortium were obtained from Lamy-Besnier^43^. These prophages belong to 7/12 bacterial strains in the OMM^12^ consortium. Additionally, several of these prophages have been induced *in vivo*^43^. The *A. muris* phage Fontainebleau was excluded from downstream analyses since its host was not included in the strains gavaged into GF mice. Quality-controlled reads were mapped to the remaining 12 prophages using Bowtie2 (v 2.2)^85^. Using bulk metagenome reads and PropagAtE (default settings, v.1.1.0)^62^ the prophage-host ratio for *C. innocuum* phage Amboise was calculated.

### Phage metagenome analyses from Duerkop *et al*. and from Lloyd-Price *et al*. datasets

Raw sequencing reads were obtained (accession number PRJEB22710) from a mouse colitis dataset from Duerkop *et al*.^65^ and a longitudinal human IBD cohort (https://ibdmdb.org/) from Lloyd-Price *et al*.^6^ In the mouse T-cell colitis dataset, bulk metagenome and phageome reads were obtained from C57BL6/J *Rag1^−/−^* mice receiving T-cell colitis (n=3) or a saline control (n=3) (Supplemental Fig. S4A). Samples were obtained at 3 time points over the course of colitis. Paired end sequencing reads (150 bp) were generated using Illumina HiSeq 2500. Raw reads were trimmed based on sequence quality, and adapter sequences were removed using Trimmomatic (v 0.33)^84^. Quality-controlled sequences that aligned to human and mouse genomes were removed by Bowtie (v 2.2)^85^. Reads originating from the same mouse or individual were grouped and co-assembled into scaffolds using Megahit (v 1.2.9)^88^. Scaffolds within each co-assembly < 3 kb in length were removed^22^ and classified as phage by VIBRANT (v 1.2.1) using the-virome flag^89^. Scaffolds were dereplicated with Blastn (v 2.14), using thresholds of an average nucleotide identity of 95% over 85% alignment fraction^66^. Scaffolds with a completeness ≥ 50%, as determined by CheckV^90^ (v 1.5) were considered as distinct vOTUs^66^ and kept for downstream analyses. Bacphlip (v 0.9.6)^69^ was used to predict phage replication cycles, classifying each vOTU as either “temperate” or “virulent”. iPHoP (v 1.0.0)^67^, which integrates several bioinformatic phage-host matching strategies, was used to predict the bacterial host of all vOTUs. Putative assignments with the highest confidence score were selected as the predicted bacterial host. We considered PHFs to be a distinct taxonomic unit, referring to the group of vOTUs that are predicted by iPHoP to infect the same bacterial family^68^. Raw reads were mapped to the vOTUs using Bowtie2^85^ and were considered present in a given sample based on mapping thresholds proposed by Stockdale^41^.

### Quantification and statistical analyses

All statistical tests indicated in figure legends were performed in R (v 4.0.5) or in GraphPad Prism (v 8.4.3). For *in vitro* prophage induction assays, AUCs were calculated using GraphPad Prism (v 8.4.3). Heatmap and visualizations of VLP fold-change were generated using ggplot2 (v 3.4.4). All relative abundance, PCoA, and PCA plots were generated using microViz (v 0.10.0.10). Aitchison’s distance, BC dissimilarity and PERMANOVA were calculated using Vegan (v 2.5.7)^91^. DESeq2 (v 1.3.0)^92^ was used to perform differential abundance analyses. In the DSS and T-cell models of colitis datasets, the standard DESeq2 workflow was performed, while controlling for baseline time points and repeated sampling of mouse cages to identify phage and bacterial taxa that were enriched during peak colitis. In the HMP-2 dataset, DESeq2 was used to identify differentially abundant PHFs based on dysbiosis status, accounting for the paired nature of the samples. Only PHFs that were present in more than 10% of samples were included in differential abundance analyses. Taxa were considered differentially abundant if the log_2_ fold change was ≤ −1 or ≥ 1 and the adjusted *p*-value ≤ 0.05. A more strict *p-*value cutoff (≤ 0.001) was used in the HMP-2 dataset to minimize false positive hits given the high number of samples and taxa.

### Key resource table

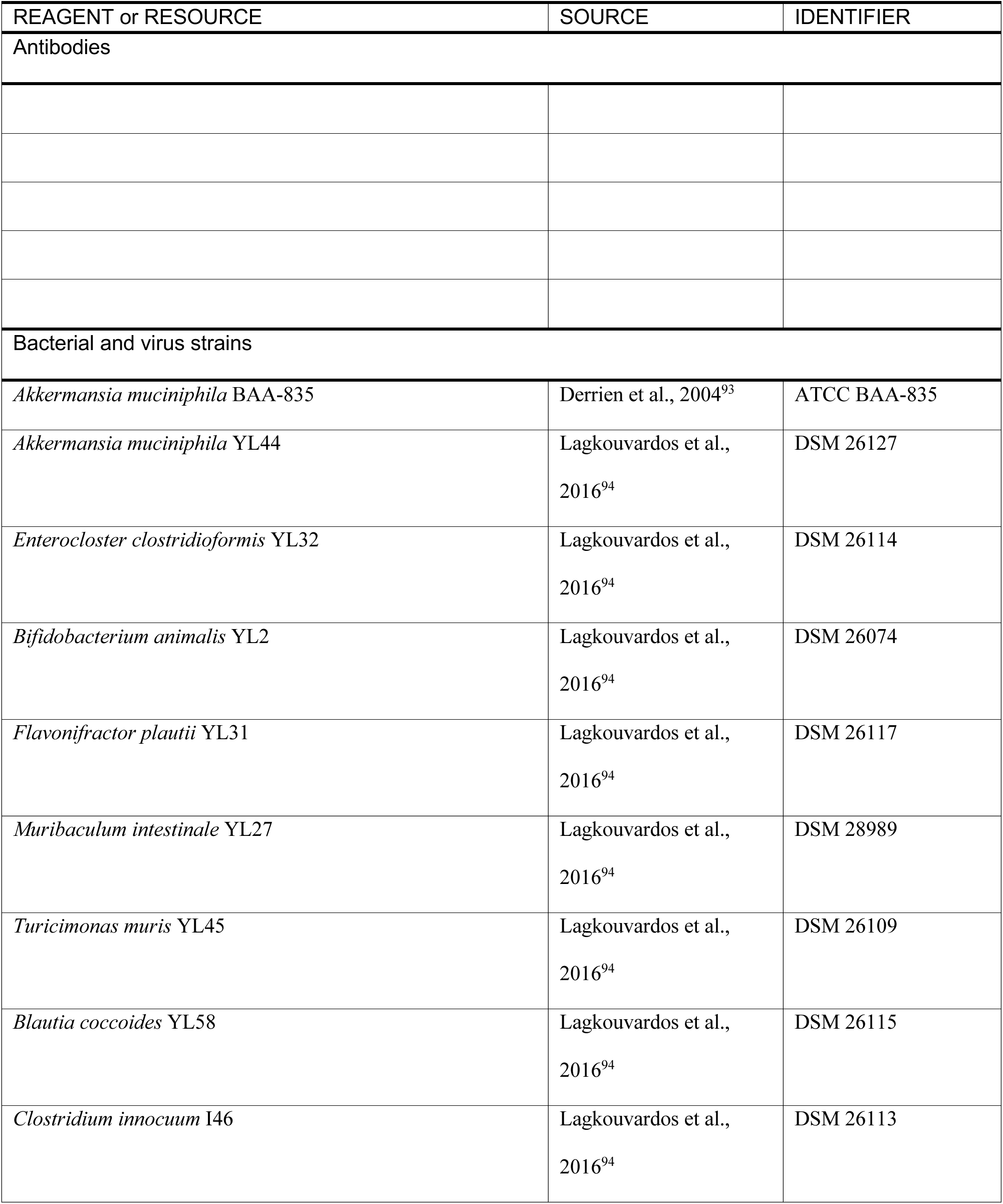
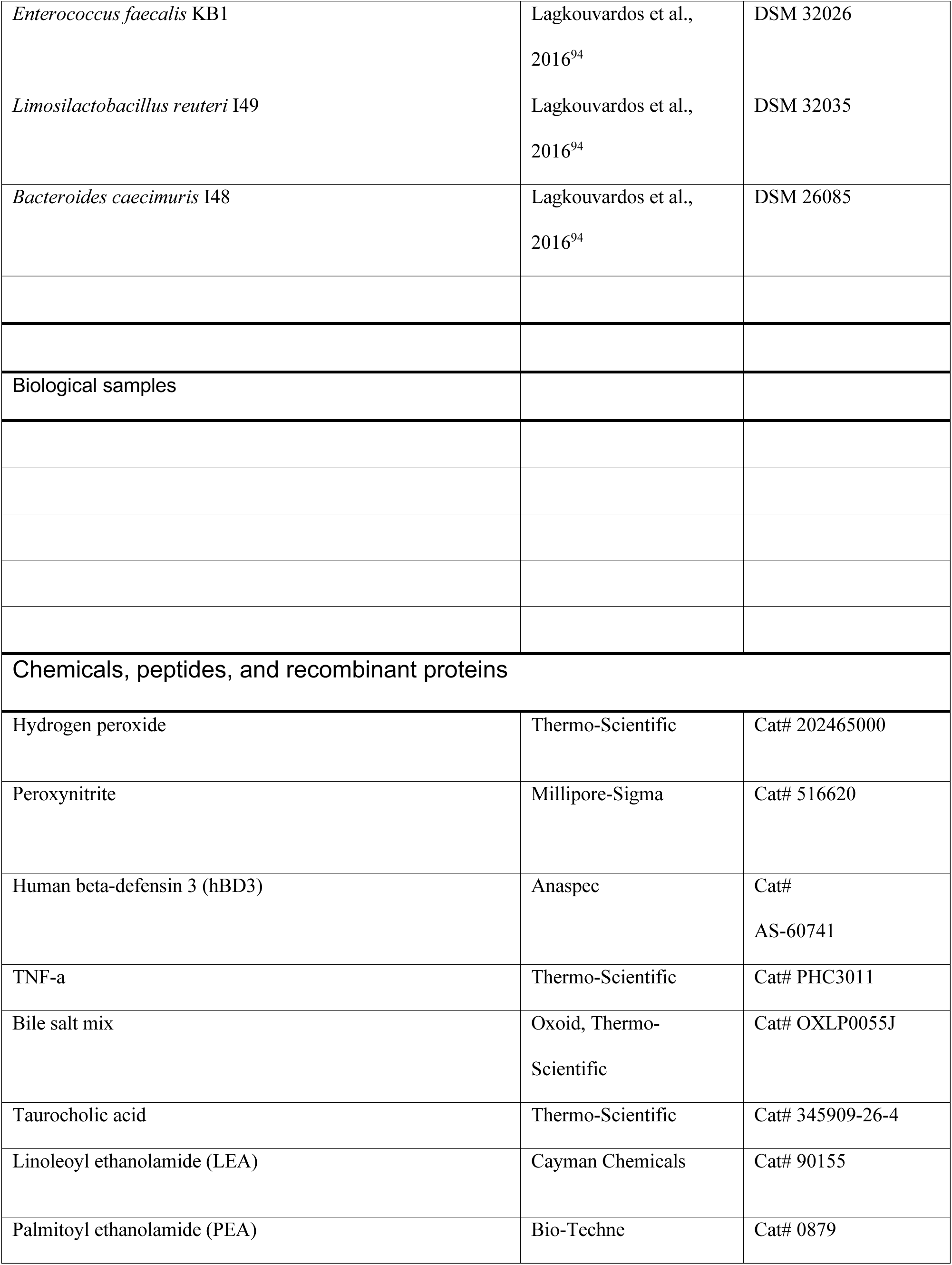
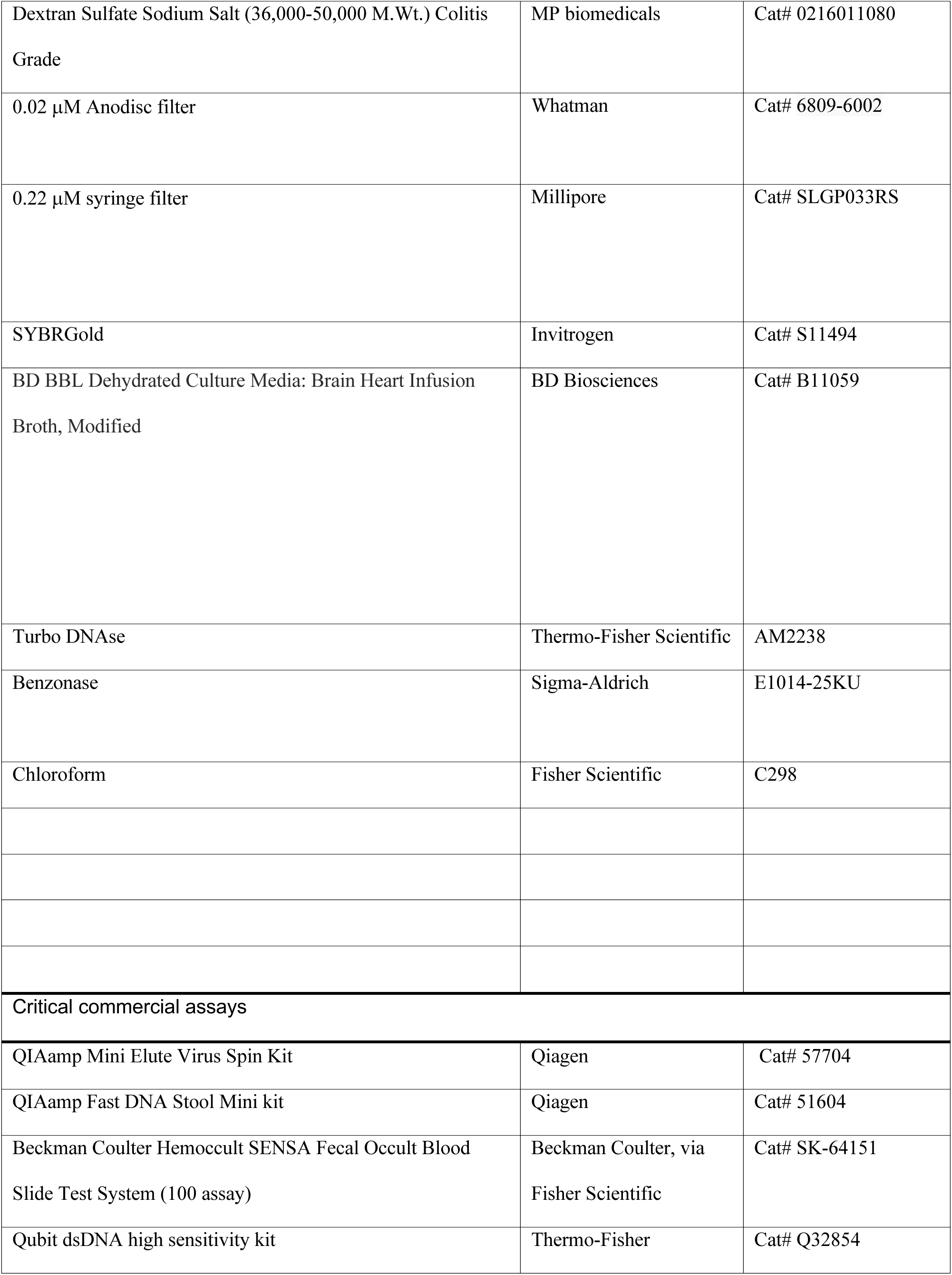
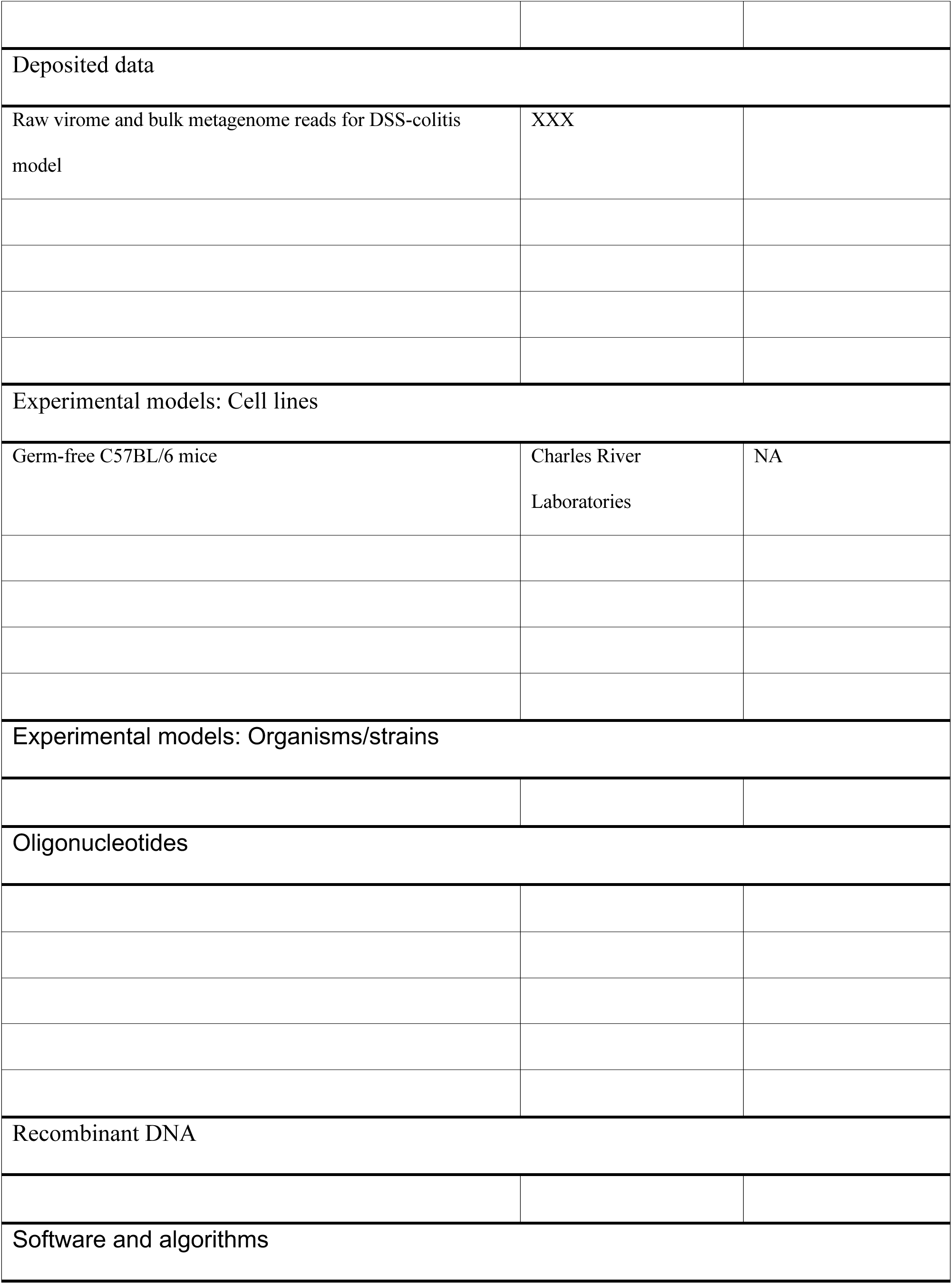
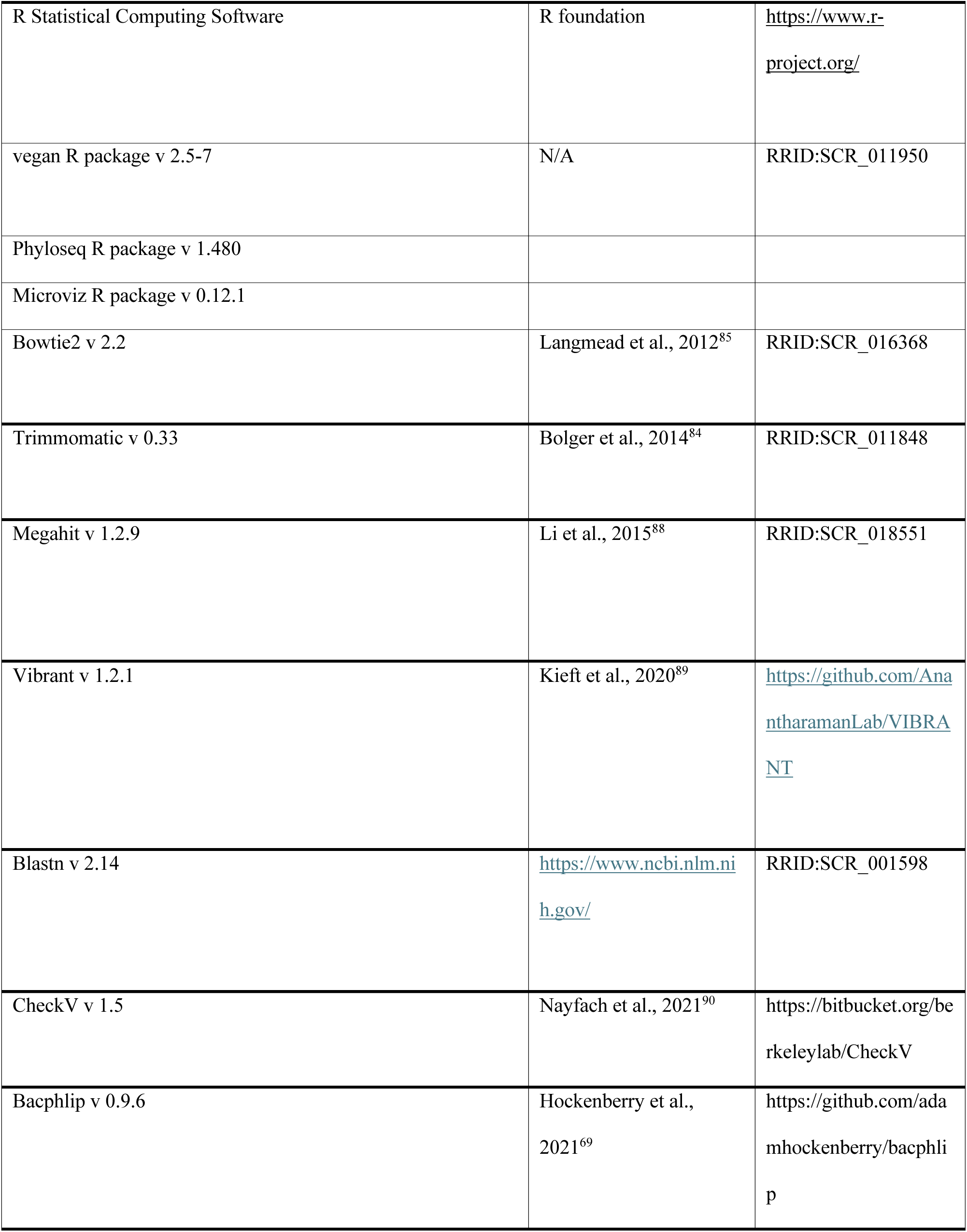
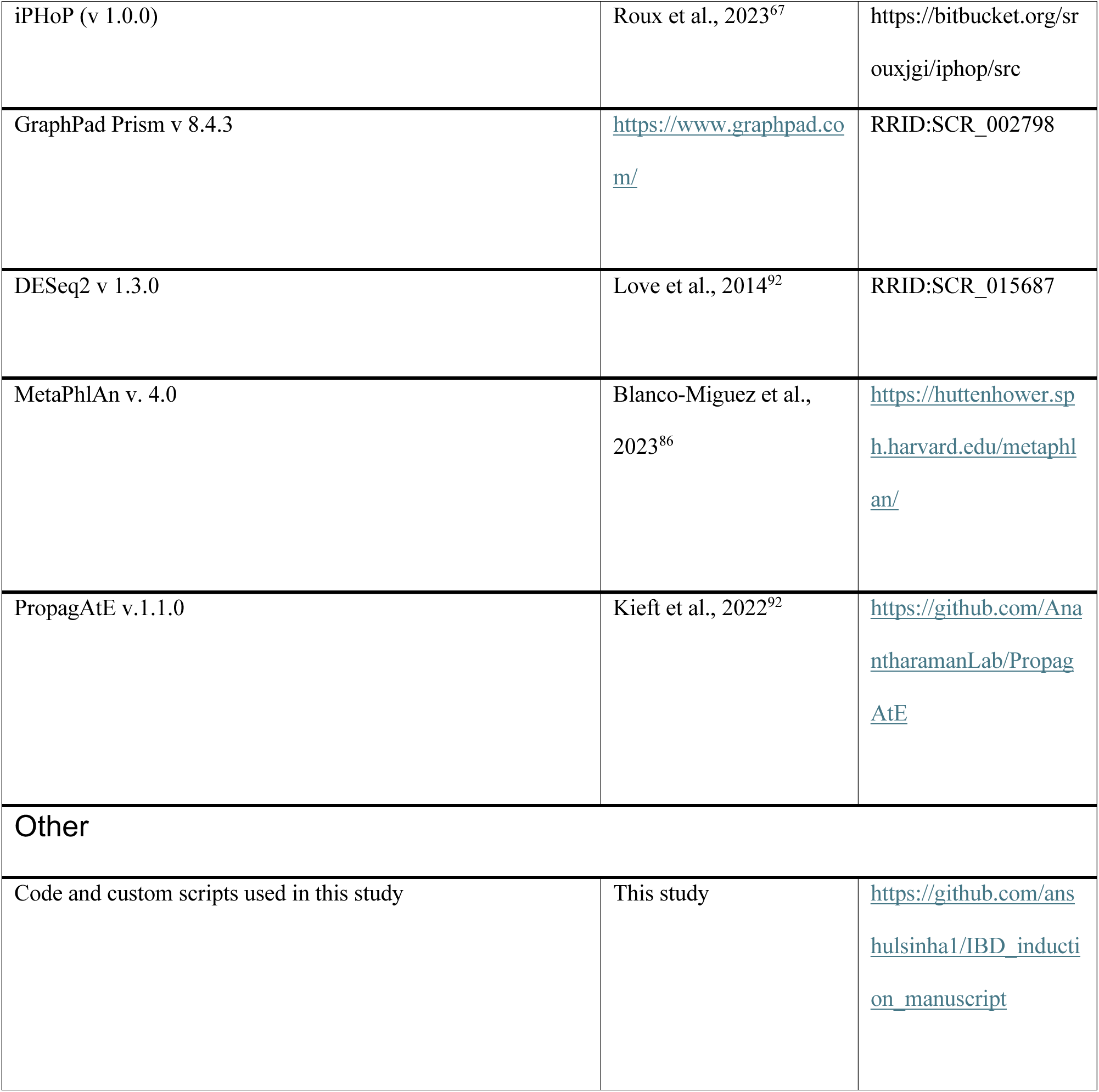

