## Supplemental Figs. 1-5, Supplemental Table 1 for "Prophage induction contributes to alterations in the gut phageome during intestinal inflammation"

### Supplemental Figures and Tables

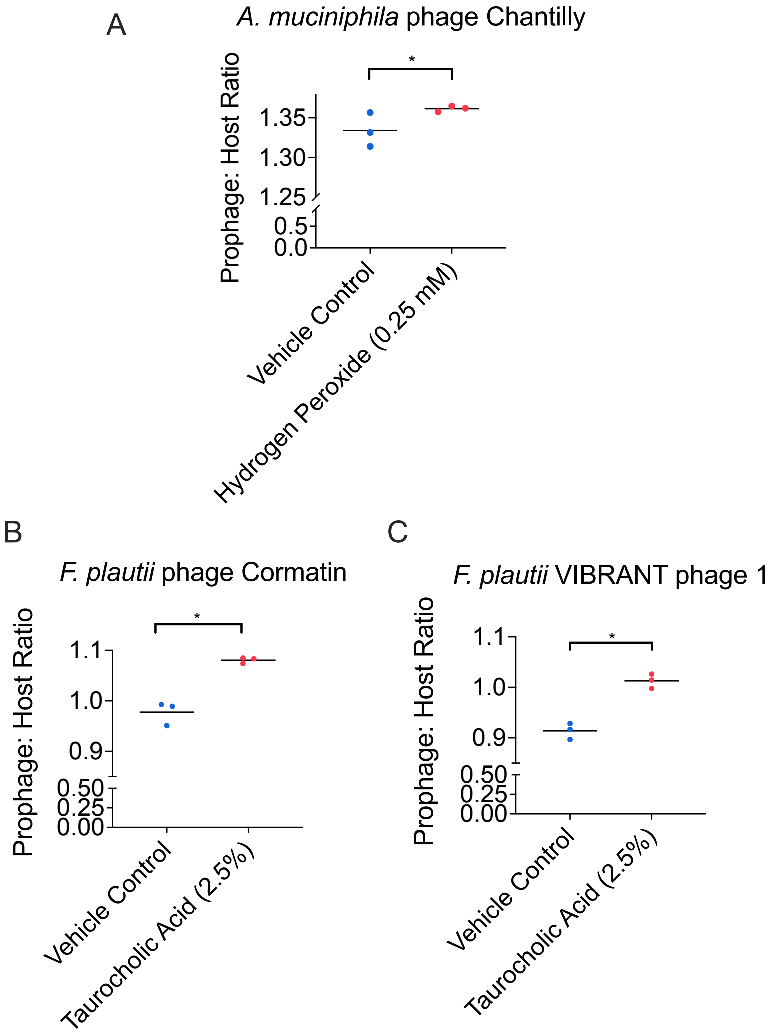


**Supplemental Fig. S1 Prophage:host ratio-based validation of prophage induction. Related to Fig. 1.**

Bulk DNA was extracted from *in vitro* bacterial cultures at stationary phase grown anaerobically in triplicate with a vehicle control or (A) hydrogen peroxide (0.25 mM) with *A. muciniphila* YL44 or (B,C) *F. plautii* YL31 grown with taurocholic acid (2.5%). Prophage:host ratios were calculated using Propagate, which aligns reads to prophage and host regions. Significant differences were assessed using the Mann-Whitney test (* $p \leq 0.05$).

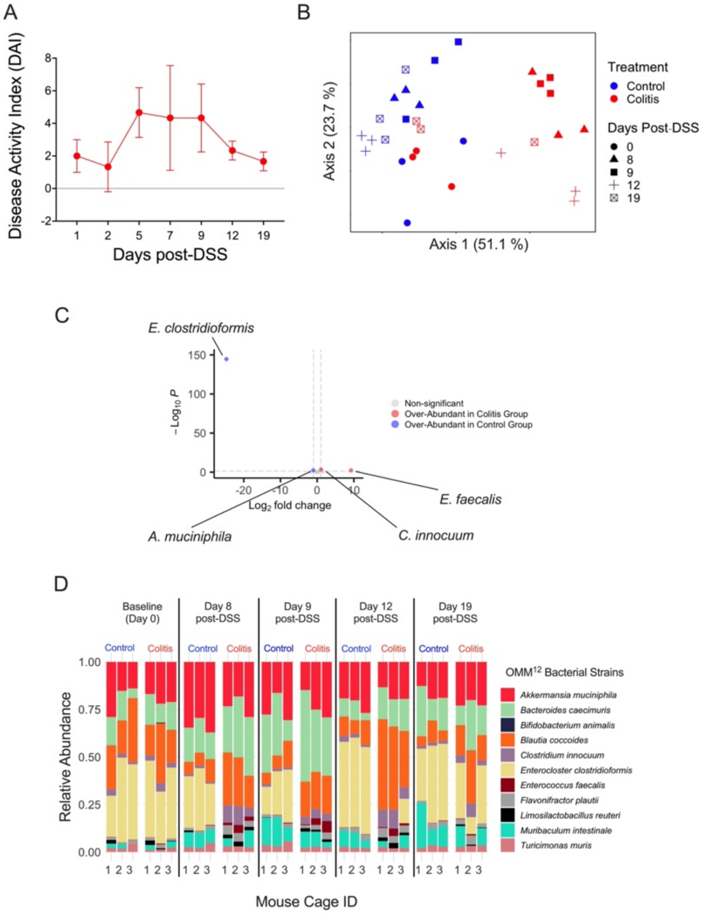


**Supplemental Fig. S2 Disease severity and bacterial community shifts during DSS-colitis in OMM$^{12}$-colonized mice. Related to Fig. 2.**

A) DAI was calculated in mice experiencing 2% DSS-colitis as the sum of weight loss, stool consistency, and hemoccult scores (see methods). Fecal samples were pooled from the same cage to measure stool consistency and the presence of fecal blood. The mean weight loss score was taken from each cage. Error bars = SD. No significant differences were found between any two time points as determined by the Friedman test ($p \leq 0.05$). n=3. B-D) Bulk community DNA was extracted from mouse feces. B) PCoA on Bray-Curtis (BC) dissimilarity of the OMM$^{12}$ bacterial taxa. The colitis group diverges from the control group in all cages on days 8, 9 and 12 post-DSS (n=3 per treatment group). C) DESeq2 was used to determine differentially abundant bacterial taxa between the control and colitis group during peak colitis (days 8 and 9 post-DSS). Taxa with a $\log_2$ fold-change $\geq 1$ or $\leq -1$ and an adjusted *p*-value $\leq 0.05$ were considered differentially abundant using the Wald test. n=3 per treatment group.

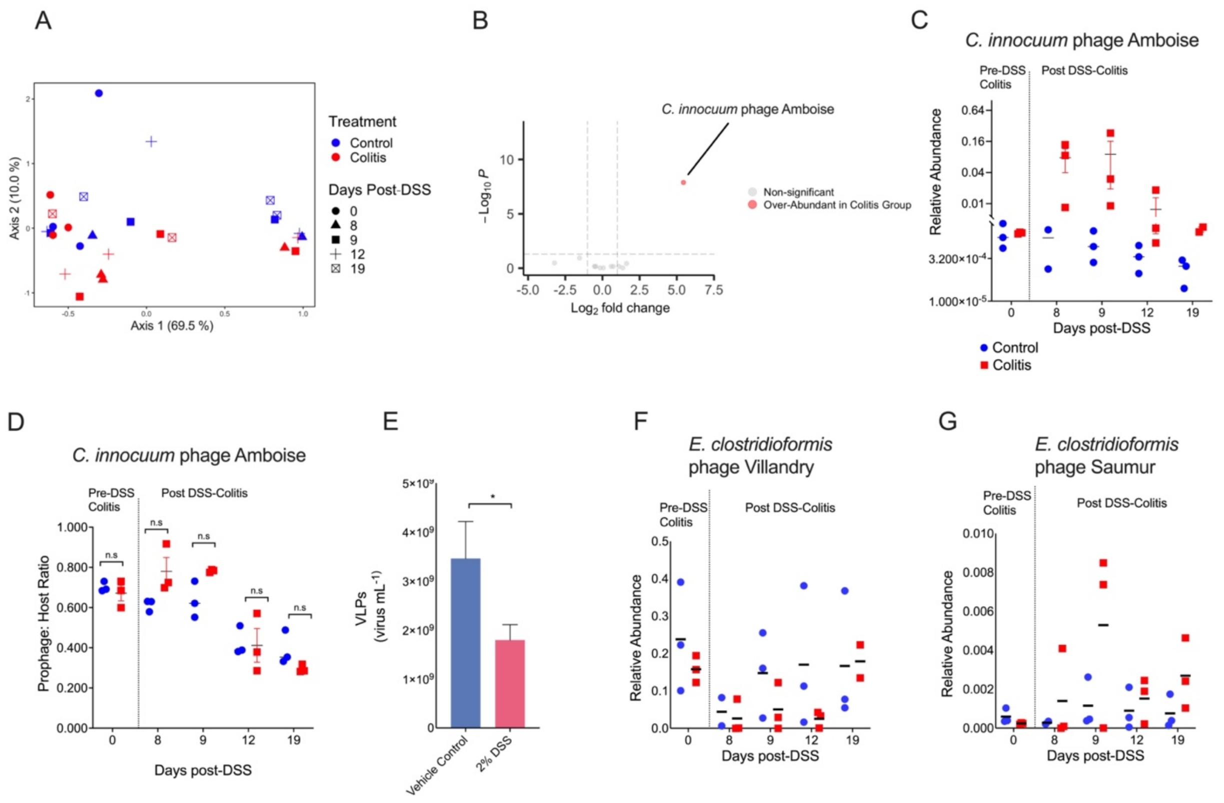


**Supplemental Fig. S3 Extracellular prophages in the OMM[12] community shift in response to DSS colitis. Related to Fig. 3.**

Fecal samples within cages were collected and pooled (n=3 per treatment group) to extract either bulk metgenome or viral DNA. Due to low biomass, viromes for 1 control sample (day 8 post-DSS) and 1 colitis group sample (19 days post-DSS) could not be obtained (n=2 per treatment group). A) PCoA on Bray-Curtis (BC) dissimilarity of extracellular OMM[12] prophages from fecal viromes. B) DESeq2 was used to determine differentially abundant OMM[12] prophages between the control and colitis group during peak colitis (days 8 and 9 post-DSS) in fecal viromes. *C. innocuum* phage Amboise was the only phage identified as differentially abundant. Taxa with a $\log_2$ fold-change $\geq 1$ or $\leq -1$ and an adjusted $p$-value $\leq 0.05$ were considered differentially abundant using DESeq2 and the Wald test. n=3 per treatment group. C) Relative abundance of *C. innocuum* phage Amboise in the fecal viromes of OMM[12]-colonized mice. Mean relative abundance is shown as a horizontal line. The y-axis is $\log_{10}$ scaled. D) Prophage-to-host ratio of *C. innocuum* phage Amboise from bulk metagenomes. PropagAtE[60] was used to assess the prophage-to-host ratio of phage Amboise. Read coverage within the genomic coordinates of phage Amboise was compared to non-prophage containing host regions. Significance ($p \leq 0.05$) was assessed between treatment groups using a repeated measures two-way ANOVA and Sidak's multiple comparison test. Mean prophage-to-host ratio is shown as a horizontal line. (E) VLP quantification in the supernatant of *C. innocuum* grown anaerobically *in vitro* with 2% DSS or the vehicle control (n=3 per treatment group). VLPs were quantified at stationary phase using epifluorescence microscopy. Significant increases in fold-change between treatment and vehicle controls were assessed using an unpaired t-test (*$p \leq 0.05$). (F) Relative abundance of *E. clostridioformis* phage Villandry and (G) Saumur in the fecal viromes of OMM[12]-colonized mice. Mean relative abundance is shown as a horizontal line.

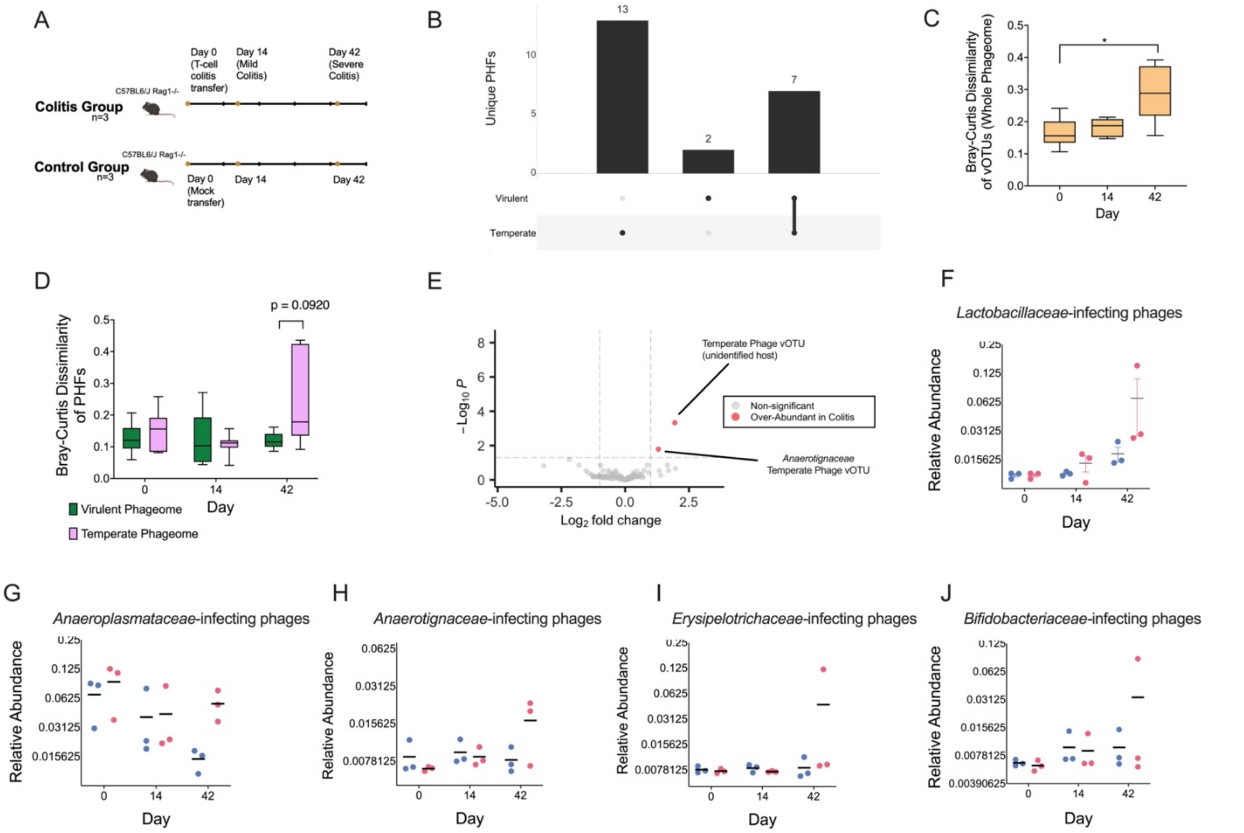


**Supplemental Fig. S4 Reanalyses of a murine T-cell colitis dataset reveals shifts to the temperate phageome. Related to Fig. 4**

A) Schematic of experimental timeline. Three C57BL6/J Rag1-/- mice received $CD4^{+}$ $CD45RB^{High}$ T cells on day 0 to induce chronic inflammation. Three C57BL6/J Rag1-/- mice received a saline control (n=3). Mice were housed individually and monitored for 6 weeks for markers of inflammation. Feces were collected on day 0, 14, and 42 for DNA extraction from VLPs. Fecal virome data were obtained from publicly available data generated by Duerkop[63]. B) Unique PHFs in the virulent and temperate phageome. The number of distinct PHFs within the virulent and temperate subsets of the whole phageome are noted at the top of the bar graphs, while the number of identified shared and unique PHFs are indicated in the UpSet plot below. A PHF was considered present if it was found in one mouse in at least one time point. C) Bray-Curtis (BC) dissimilarity of the whole free phageome at the viral OTU (vOTU) level of phage taxonomy. Significance was determined using the Friedman test (* $p \leq 0.05$). (D) BC dissimilarity between the control and colitis groups at the PHF level of phage taxonomy in the temperate and virulent subsets of the whole phageome. Significance was assessed using a repeated measures two-way ANOVA using Sidak's multiple comparison test, $p \leq 0.05$. (E) DESeq2 was used to determine differentially abundant vOTUs in the free phageome during peak colitis between the control and colitis groups in a T-cell model of colitis[90]. vOTUs with a $\log_2$ fold-change ≥ 1 or ≤ -1 and with *p*-values ≤ 0.05 were considered differentially abundant using the Wald test. (F-J) Relative abundance of PHFs over time. PHFs in the free phageome were assigned a host using iPHoP[65]. Mean relative abundance is shown by horizontal lines.

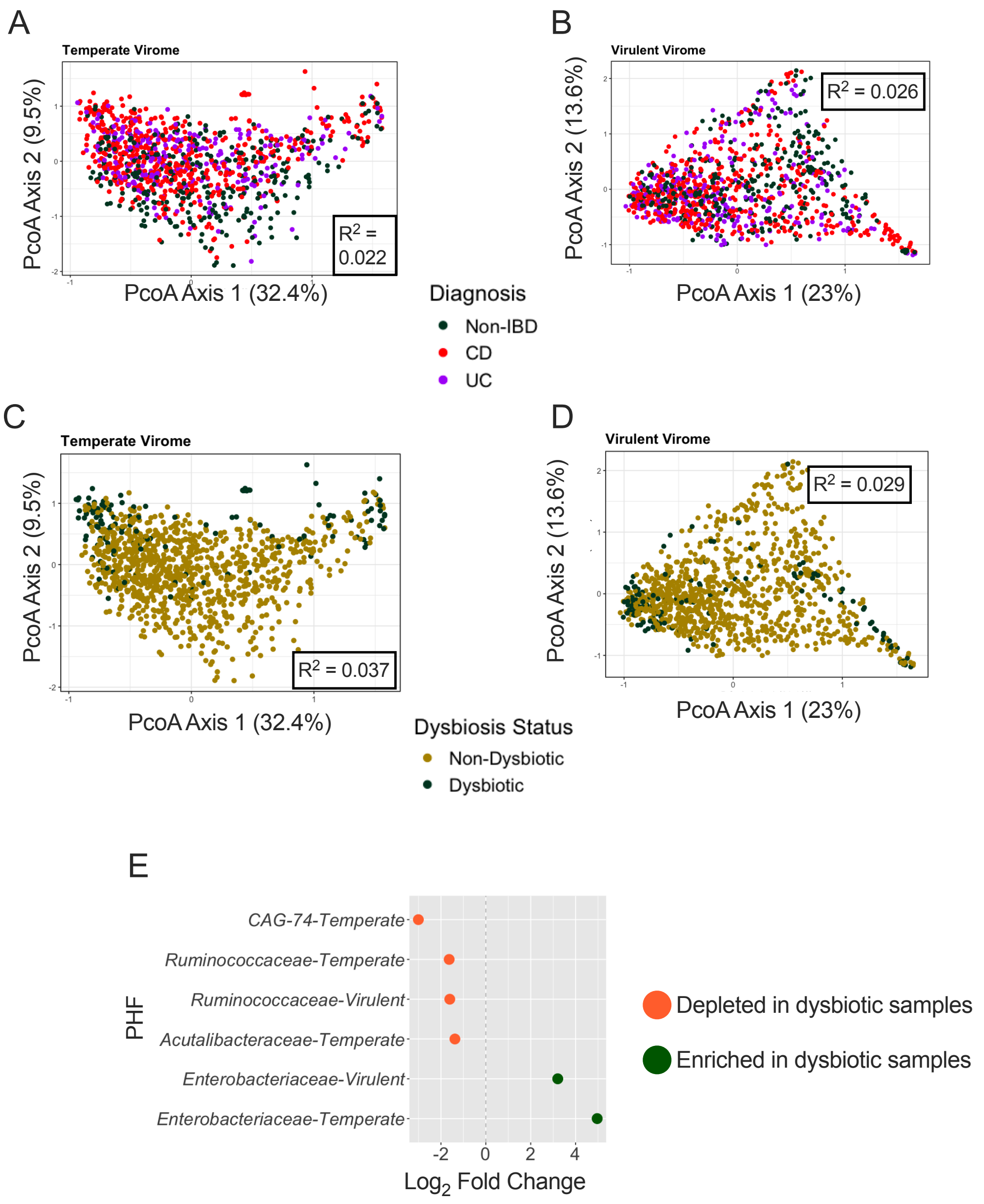


**Supplemental Fig. S5 Phageome diversity of IBD patients in temperate and virulent subsets. Related to Fig. 4**

Bulk metagenomes were obtained from publicly available data generated by Lloyd-Price[6]. IBD patients and non-IBD controls were sampled longitudinally over 1 year (n=1,595 samples from 130 individuals). Dysbiotic samples refer to samples identified by Lloyd-Price as being highly dissimilar in bacterial composition to non-IBD controls[6]. PCoA on Bray distance based on (A-B) diagnosis status or (C-D) dysbiosis status. Bray distance was calculated at the PHF level of taxonomy. n=1,093 from 115 individuals. E) PHFs were split by putative replication cycle and differential abundance was determined using DESeq2[90]. PHFs with a $\log_2$ fold change $\geq 1$ and an adjusted *p*-value $\leq 0.001$ indicate those significantly enriched in dysbiotic samples (green). PHFs with a $\log_2$ fold change $\leq -1$ and an adjusted *p* value $\leq 0.001$ indicate those significantly depleted in dysbiotic samples (orange). For differential abundance analyses, individuals were only included if they had both a dysbiotic and non-dysbiotic sample. n=487 samples from 49 individuals.

| Bacterial Strain | Phylum | Prophage Identification Method | Nucleotide Start Position | Nucleotide Stop Position | Prophage Length (bp) |
|---|---|---|---|---|---|
| *Akkermansia muciniphila* BAA-835 | Verrucomicrobiota | Prophage 1-VIBRANT | 97,910 | 104,338 | 6,429 |
| | | Prophage 2-VIBRANT | 1,870,783 | 1,901,246 | 30,464 |
| *Akkermansia muciniphila* YL44 | Verrucomicrobiota | Prophage "Chambord"-Lamy-Besnier | 712,134 | 742,810 | 30,676 |
| | | Prophage "Moulinsart"-Lamy-Besnier | 1,335,300 | 1,366,646 | 31,346 |
| | | Prophage "Chantilly"-Lamy-Besnier | 2,300,756 | 2,337,649 | 36,893 |
| *Bacteroides caecimuris* I48 | Bacteroidota | Prophage "Versailles"-Lamy-Besnier | 2,968,431 | 3,026,527 | 58,096 |
| *Flavonifractor plautii* YL31 | Bacillota | Prophage 1-VIBRANT | 459,512 | 466,007 | 6,495 |
| | | Prophage "Castelnaud"-Lamy-Besnier | 908,701 | 945,825 | 37,124 |
| | | Prophage 3-VIBRANT | 993,247 | 1,034,458 | 4,1211 |
| | | Prophage "Cormatin"-Lamy-Besnier | 1,128,657 | 1,164,096 | 35,439 |
| | | Prophage 5-VIBRANT | 2,668,007 | 2,715,122 | 47,115 |
| | | "Chenonceau"-Lamy-Besnier | 2,742,851 | 2,784,974 | 42,123 |

**Supplemental Table. 1 List of prophages detected within bacterial strains used for *in vitro* prophage induction assays. Related to Fig. 1**
Prophages were either identified previously by Lamy-Besnier et al.,[43] or bioinformatically predicted using VIBRANT[87].
